# Three-dimensional drug screen identifies HDAC inhibitors as therapeutic agents in mTORC1*-*driven lymphangioleiomyomatosis

**DOI:** 10.1101/2021.07.03.451004

**Authors:** Adam Pietrobon, Julien Yockell-Lelièvre, Nicole Melong, Laura J. Smith, Sean P. Delaney, Nadine Azzam, Chang Xue, Nishanth Merwin, Eric Lian, Alberto Camacho-Magallanes, Carole Doré, Gabriel Musso, Lisa M. Julian, Arnold S. Kristof, Roger Y. Tam, Jason N. Berman, Molly S. Shoichet, William L. Stanford

## Abstract

Lymphangioleiomyomatosis (LAM) is a rare disease involving cystic lung destruction by invasive LAM cells. These cells harbor loss-of-function mutations in *TSC2*, conferring constitutive mTORC1 signaling. Rapamycin is the only clinically approved disease-modifying treatment, but its action is cytostatic and disease progresses upon its withdrawal. There is a critical need to identify novel agents that prevent the invasive phenotype and/or eradicate the neoplastic LAM cells. Here, we employed novel cellular and extracellular models to screen for candidate therapeutics in a physiologically relevant setting. We observed that lung-mimetic hydrogel culture of pluripotent stem cell-derived diseased cells more faithfully recapitulates human LAM biology compared to conventional culture on two-dimensional tissue culture plastic. Leveraging our culture system, we conducted a three-dimensional drug screen using a custom 800-compound library, tracking cytotoxicity and invasion modulation phenotypes at the single cell level. We identified histone deacetylase (HDAC) inhibitors as a group of anti-invasive agents that are also selectively cytotoxic towards *TSC2^-/-^* cells. Unexpectedly, we observed that next generation ATP-competitive mTORC1/2 inhibitors potentiate invasion. We determined anti-invasive effects of HDAC inhibitors to be independent of genotype, while selective cell death is mTORC1-dependent and mediated by apoptosis. Drug performance was subsequently evaluated at the single cell level in zebrafish xenografts. We observed consistent therapeutic efficacy *in vivo* at equivalent concentrations to those used *in vitro,* substantiating HDAC inhibitors as potential therapeutic candidates for pursuit in patients with LAM.

**One Sentence Summary:** We performed a drug screen in 3D and discovered HDAC inhibitors exhibit therapeutic efficacy in models of the lung disease lymphangioleiomyomatosis.

## INTRODUCTION

Lymphangioleiomyomatosis (LAM) is a cystic lung disease predominately affecting women, at a prevalence of 1 to 10 per million (*1*). LAM can occur sporadically or in association with the multisystem tumor-forming disorder, Tuberous Sclerosis Complex (TSC) (*2*). The pulmonary histopathology is characterized by microscopic nodules consisting of immature smooth muscle-like cells that express markers of neural crest lineages (*3*). These invading cells digest the lung parenchyma forming cystic lesions that lead to progressive respiratory decline and fatality if untreated (*4–6*). The molecular etiology of LAM involves loss-of-function mutations in the endogenous mTORC1 suppressor *TSC2*, thereby inducing hyperactivation of mTORC1 anabolic and tumorigenic signalling (*7*). The allosteric mTORC1-inhibitor rapamycin (clinically, sirolimus) slows disease progression and improves symptomatology (*8–11*). While clinical approval of rapamycin by the FDA in 2015 has led to a dramatic new frontier in the LAM therapeutic landscape, significant limitations exist. A subset of patients do not respond to treatment, and rapamycin is invariably cytostatic, with rapid disease progression upon treatment withdrawal (*11, 12*). There is a critical need to discover novel treatment strategies that can eradicate LAM cells.

A key step in the pathway to therapeutic development is the effective modelling of disease characteristics. In this domain, LAM has remained a challenge. Cultures of cells derived from human pulmonary LAM lesions grow as a heterogeneous mixture with rapid exhaustion of *TSC2^-/-^* cells, prohibiting the establishment of clonal primary cell lines (*13*). While a genome engineering strategy would seem straightforward for this monogenic disease, the cell-of-origin of LAM remains unknown, begging the question of which cell type to engineer. While we have demonstrated that *TSC2^-/-^* human pluripotent stem cell-derived neural crest cells model several phenotypic features of LAM (*14*), neural crest cells consist of a diverse and plastic population that are not readily scalable for drug screening purposes. Animal models of LAM have been comparably challenging to establish, and none to date have recapitulated pathognomonic features such as histological premelanosome protein (PMEL) positivity and concomitant elevated serum levels of vascular endothelial growth factor D (VEGF-D) (*15*).

An emerging consideration in disease modelling is the contribution of the extracellular matrix (ECM) to disease biology. Water-swollen networks of polymers termed hydrogels have arisen as effective tools for mimicking salient elements of the native ECM while exhibiting mechanics similar to many soft tissues (*16*). Hydrogels can be broadly classified as either natural, synthetic, or hybrid materials. One such hybrid scaffold is hyaluronic acid, a naturally-sourced material that can be readily modified to independently tune ECM features of interest, such as elasticity, stiffness, and viscosity (*17*). A viscoelastic hydrogel with a derivatized hyaluronic acid backbone has been shown to permit the study of invasive properties of LAM cellular models in three-dimensional culture (*18*). Importantly, three-dimensional culture systems have been demonstrated as more predictive of *in vivo* drug responses compared to conventional culture on two-dimensional plastic (*19, 20*).

In recent years, there has been a resurgence of interest in phenotype-based screens for drug discovery compared to target-based approaches (*21*). An analysis of therapeutics approved between 1999 and 2008 revealed that 62% first-in-class drugs were discovered by phenotype-based screens, despite the fact that such screens represented only a small subset of the overall total (*22*). The apparent superiority of phenotype-based approaches may in part arise from the ability to identify compounds which exhibit a therapeutic effect by modulating multiple targets simultaneously (*21*). In addition, phenotypic drug screens can be multiplexed with counter-screening, ensuring candidate therapeutics do not also confer undesirable side-effects, such as physiological toxicity. In the context of LAM, a monogenetic disease, this counter-screening takes shape by directly comparing *TSC2^-/-^* cells against matched wild type (WT) controls.

Here, we analyze a novel hydrogel culture system of pluripotent stem cell-derived models, and observe the cell type employed, genotype, and culture substrate all contribute to modelling features of LAM. We performed a three-dimensional drug screen, tracking cytotoxicity and invasion modulation phenotypes at the single cell level. We identified histone deacetylase (HDAC) inhibitors as anti-invasive and selectively cytotoxic towards *TSC2^-/-^* cells. Importantly, we observed consistent therapeutic efficacy upon xenotransplantation of human cell models into zebrafish larvae, highlighting HDAC inhibitors as potential therapeutic candidates for pursuit in patients.

## RESULTS

### Stem cell-derived models exhibit features of LAM, independent of genotype

As pulmonary LAM cells are not amenable to expansion upon lesion explant, (*13*) we established primary cell lines by *in vivo* differentiation of human pluripotent stem cells (hPSCs), as previously described (*23*). Briefly, hPSCs were injected into NOD.Cg-*Prkdc^scid^ Il2rg^tm1Wjl^*/SzJ (NSG) immunodeficient mice to form teratomas, which were explanted and expanded in smooth muscle-cell enriching conditions (Fig. S1A). We used a previously reported isogenic pair of female mCherry^+^ WT and genome-engineered *TSC2*^-/-^ hPSCs (*14*). Cell cultures exhibit a predominately spindle cell morphology and express α-smooth muscle actin (ACTA2) protein in all isolated cells (Fig. 1A, Fig. S1B). Further, immunofluorescence analysis identified a small fraction of PMEL^+^ cells (∼0.13%), a hallmark marker of pulmonary LAM (Fig. 1A, Fig. S1C). The high fraction of ACTA2^+^ and low fraction of PMEL^+^ cells in culture is consistent with the relative abundance of these markers in heterogenous human LAM lesions (*3*). Notably, the percentage of PMEL^+^ and ACTA2^+^ cells did not vary between WT and *TSC2^-/-^* (Fig. S1B-C). Secreted VEGF-D, a critical biochemical biomarker used in the diagnosis of LAM, was detected in the supernatant of both WT and *TSC2^-/-^* cultures and was insensitive to acute rapamycin treatment (Fig. 1B). Together, these data suggest the cell models employed exhibit features of LAM as a product of the cell type isolated, independent of genotype.

**Fig. 1.**
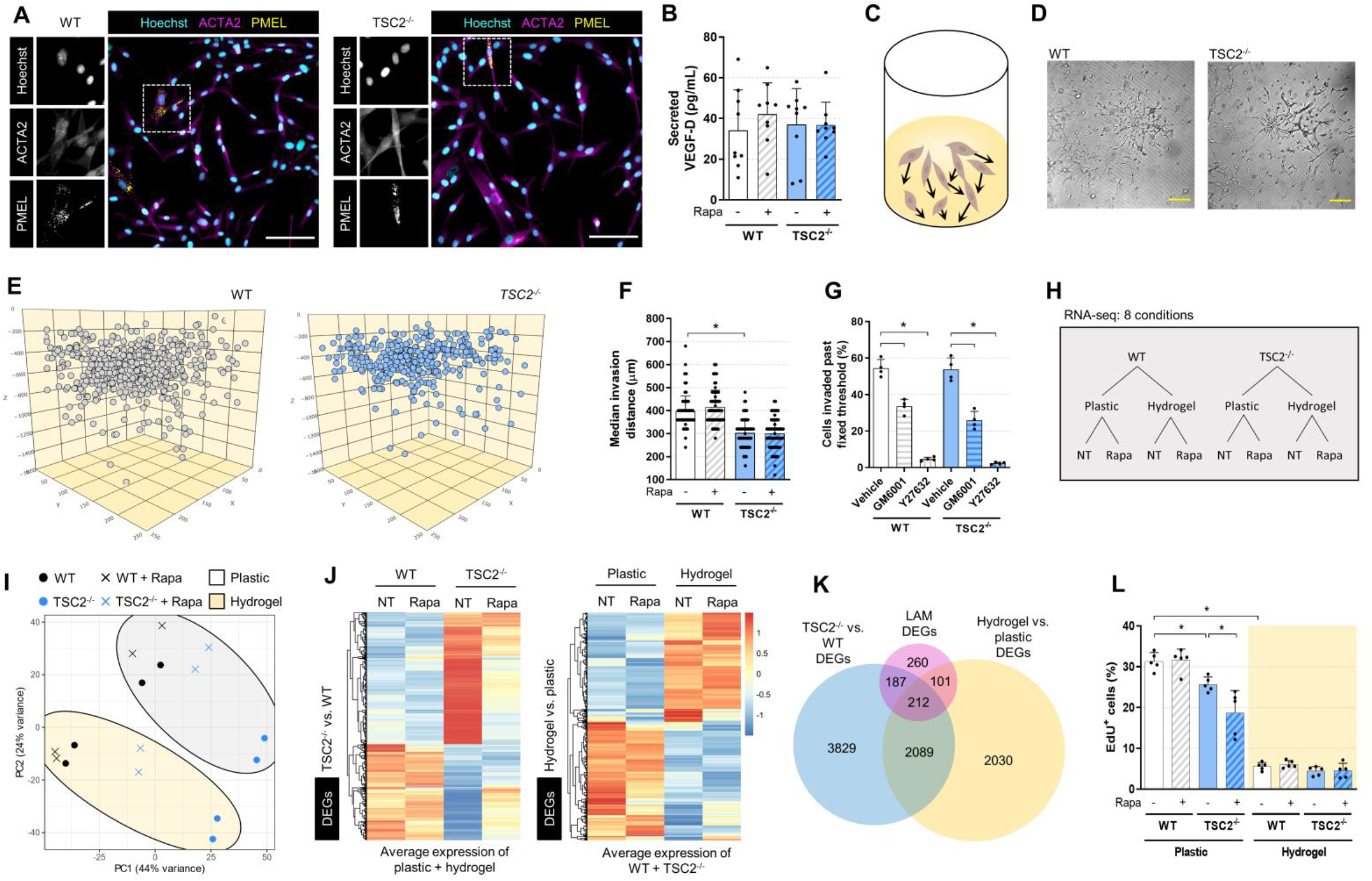
Hydrogel culture of stem cell-derived disease models exhibits features of LAM. (**A**) Representative immunofluorescence images of *WT* and *TSC2^-/-^* cells. Inset showing punctate PMEL and fibril ACTA2 staining. Scale bars of 100µM. (**B**) VEGF-D secreted into conditioned media measured by ELISA, following 16hr incubation in serum-free media ± 20nM rapamycin (mean ± SD; * = p < 0.05 by Student t-test; n = 9-10). (**C-E**) Visualization of LAM cell invasion after three days in hydrogel culture, as (**C**) a schematic, (**D**) brightfield image of single Z plane, scale bars of 250µm, and (**E**) computational reconstruction of cellular spatial positions. (**F**) Median invasion distance of cellular populations plated on the hydrogel and cultured for three days ± 20nM rapamycin (mean ± SD; * = p < 0.05 by Student t-test; n = 124). (**G**) Percentage of cells invaded past fixed threshold set by median invasion distance of genotype-matched vehicle control. Cells were cultured and treated for three days (10µM GM6001, a pan-MMP inhibitor, and 20µM Y27632, a ROCK inhibitor, mean ± SD; * = p < 0.05 by Student t-test; n = 4). (**H**) Schematic of the sample conditions tested in the bulk RNA-seq experiment. NT = no treatment, Rapa = rapamycin treatment (20nM, 72 hrs). (**I**) Principal components analysis (PCA) of bulk RNA-seq samples. (**J**) Heatmap and hierarchal clustering of differentially expressed genes (DEGs) between *TSC2^-/^*^-^ and WT samples, and between hydrogel and plastic samples, while controlling for the reciprocal covariate. Left panel: transcript expression for plastic and hydrogel cultures were averaged. Right panel: transcript expression for WT and *TSC2^-/-^* samples were averaged. DEG analysis was performed with no treatment samples; genes noted as differentially expressed if FDR < 0.05 and |log2FC| > 1. (**K**) Overlap in DEG between genotype and ECM gene lists and LAM cell signature gene list (*24*). Genes noted as DE if FDR < 0.05. (**L**) Percentage of EdU^+^ (proliferating) cells from 3-hour pulse (5µM), after three days cultured on plastic or hydrogel ± 20nM rapamycin (mean ± SD; * = p < 0.05 by Student t-test; n = 5).

### Three-dimensional hydrogel culture enables study of the LAM invasive phenotype at single cell resolution

We next sought to model the pulmonary invasive phenotype of LAM cells by adapting a lung-mimetic hydrogel culture system (*18*). The hydrogel is synthesized by crosslinking hyaluronic acid strands with matrix metalloprotease (MMP)-cleavable peptides, while embedding vitronectin peptides and methylcellulose to increase cell adhesion and matrix plasticity, respectively. Cells are plated on top of the synthesized hydrogel and actively invade through the material (Fig. 1C-D, Supplementary Movies 1-2). By staining with a nuclear dye and acquiring multiplanar images through the optically clear hydrogel, we identify every cell in XYZ planes and compute invasion distances at single cell resolution (Fig. S1D).

We observed all cells from both WT and *TSC2^-/-^* cultures to invade through the hydrogel, albeit at variable distances (Fig. 1E). On average, WT cultures invaded further than *TSC2^-/-^* in a manner insensitive to acute rapamycin treatment (Fig. 1F). We posited that differing invasion distances of cells in the same culture reflect a cell autonomous property, rather than a reflection of stochasticity. To test this, we isolated and expanded clones from WT and *TSC2^-/-^* bulk cultures and subjected these clones to hydrogel culture. We observed a subset of clones with dramatically high invasion speeds, and likewise, a subset with slow invasion speeds (Fig. S1E). These data suggest differential cell autonomous capacities for invasion in putative heterogenous cultures. Finally, we investigated modes of invasion employed by LAM cell models in this hydrogel system. Similar to previous findings (*18*), we observed a decrease in invasion upon treatment with the pan-MMP inhibitor GM6001 or the Rho-kinase (ROCK) inhibitor Y27632, indicating both protease-dependent and independent modes of invasion employed (Fig. 1G).

### Loss of *TSC2* and hydrogel culture both confer transcriptomic features of LAM

To profile our cell culture system more comprehensively, we conducted bulk RNA-seq of WT and *TSC2^-/-^* cells, in the presence or absence of rapamycin, and in both plastic and hydrogel culture, for a total of 8 sample conditions (Fig. 1H). Principal components analysis (PCA) revealed sample genotype to be driving the primary axis of variation, and culture substrate to be driving the secondary axis of variation (Fig. 1I). Rapamycin treatment induced a substantial global transcriptomic change in the *TSC2^-/-^* cells, inducing a profile more similar to WT cells (Fig. 1I).

We conducted differential expression analysis comparing across genotype (*TSC2^-/-^* vs. WT) and culture substrate (hydrogel vs. plastic) in untreated samples, while holding the reciprocal covariate constant. At a false discovery rate (FDR) < 0.05, we identified 6,317 differentially expressed genes (DEGs, 1,793 with |log_2_FC| > 1) between WT and *TSC2^-/-^*, and 4,432 DEGs (771 with |log_2_FC| > 1) between plastic and hydrogel (Table S1A-B, Fig. S1F-G). While exhibiting some overlap, these DEG lists were largely distinct (Fig. S1H). We found 78.8% of the DEGs distinguishing genotype to be reversed by rapamycin treatment, suggesting mTORC1-dependency (Fig. 1J, left panel). In contrast, the expression of DEGs distinguishing plastic versus hydrogel cultures remained largely unchanged in the presence of rapamycin (Fig. 1J, right panel). We next examined the overlap of these DEGs with a recently published LAM gene signature derived from single cell RNA-seq profiling of primary lesions (*24*). We observed that both DEG lists overlap substantially (65.8% of the total 760 LAM genes) and share both common and distinct genes with the LAM gene signature (Fig. 1K).

To glean further biological insight, we conducted GO term enrichment (Table S2A-B, Fig. S1I-J). Both DEG lists ranked “extracellular matrix organization” as most highly enriched, which is also the top enriched term in a primary LAM lesion gene signature list (*24*). The DEGs distinguishing genotype were also enriched in many terms related to development, similar to primary LAM lesions (*24*). Interestingly, the DEGs distinguishing culture substrates were largely enriched in terms related to proliferation (Table S1B, Fig. S1J). LAM is an indolent disease which progresses at a slow pace relative to other invasive diseases; only a small fraction of cells actively proliferative in primary LAM lesions (*3*). On plastic, we found LAM cell models proliferated rapidly, with ∼30% of cells incorporating EdU after a short 3-hour pulse (Fig. 1L). In contrast, *TSC2^-/-^* cells proliferated at a slightly slower pace, consistent with previous studies of loss of *TSC2* in primary cells (*25*). Acute rapamycin treatment reduced proliferation of *TSC2^-/-^* cells but did not have a detectable effect on WT cultures. However, subjecting cells to hydrogel culture caused a dramatic decrease in cell proliferation (Fig. 1L), likely reflective of the proliferation-invasion dichotomy (*26*). Together, these data suggest both genotype (loss of *TSC2*) and culture substrate (3D hydrogel) induce transcriptomic landscapes which model LAM features.

### Hydrogel culture potentiates differential mTORC1-signalling between WT and *TSC2^-/-^* cells

mTORC1 hyperactivation is a hallmark feature of primary LAM lesions compared to normal adjacent WT tissue. To assess mTORC1 signalling status, we performed a low input western blot, probing for downstream mTORC1 effectors pS6RP and p4E-BP1. Culturing cells on plastic (2D) showed marginal differences in mTORC1 signalling between WT and *TSC2^-/-^* cells, and both cell types demonstrated activation of mTORC1 above rapamycin-treated levels (Fig. 2A). Remarkably, culturing on hydrogel potentiated a dramatic difference in mTORC1 signalling, with WT cells downregulating activity to rapamycin-treated levels and *TSC2^-/-^* cells upregulating signalling above levels seen on plastic alone. This is consistent with the PCA of transcriptomic landscapes, whereby WT untreated and WT rapamycin-treated samples from hydrogel culture cluster slightly more closely compared to plastic culture (Fig. 1I). To corroborate these findings at the single cell level, we examined mTORC1 signalling by immunofluorescence (Fig. 2B, S2A). While a small difference in mTORC1 signalling was observed between WT and *TSC2^-/-^* cells cultured on plastic, this difference was potentiated in 3D hydrogel culture. Importantly, mTORC1 signalling in WT cells was seen to mirror rapamycin-treated levels only when cultured on hydrogel.

**Fig. 2.**
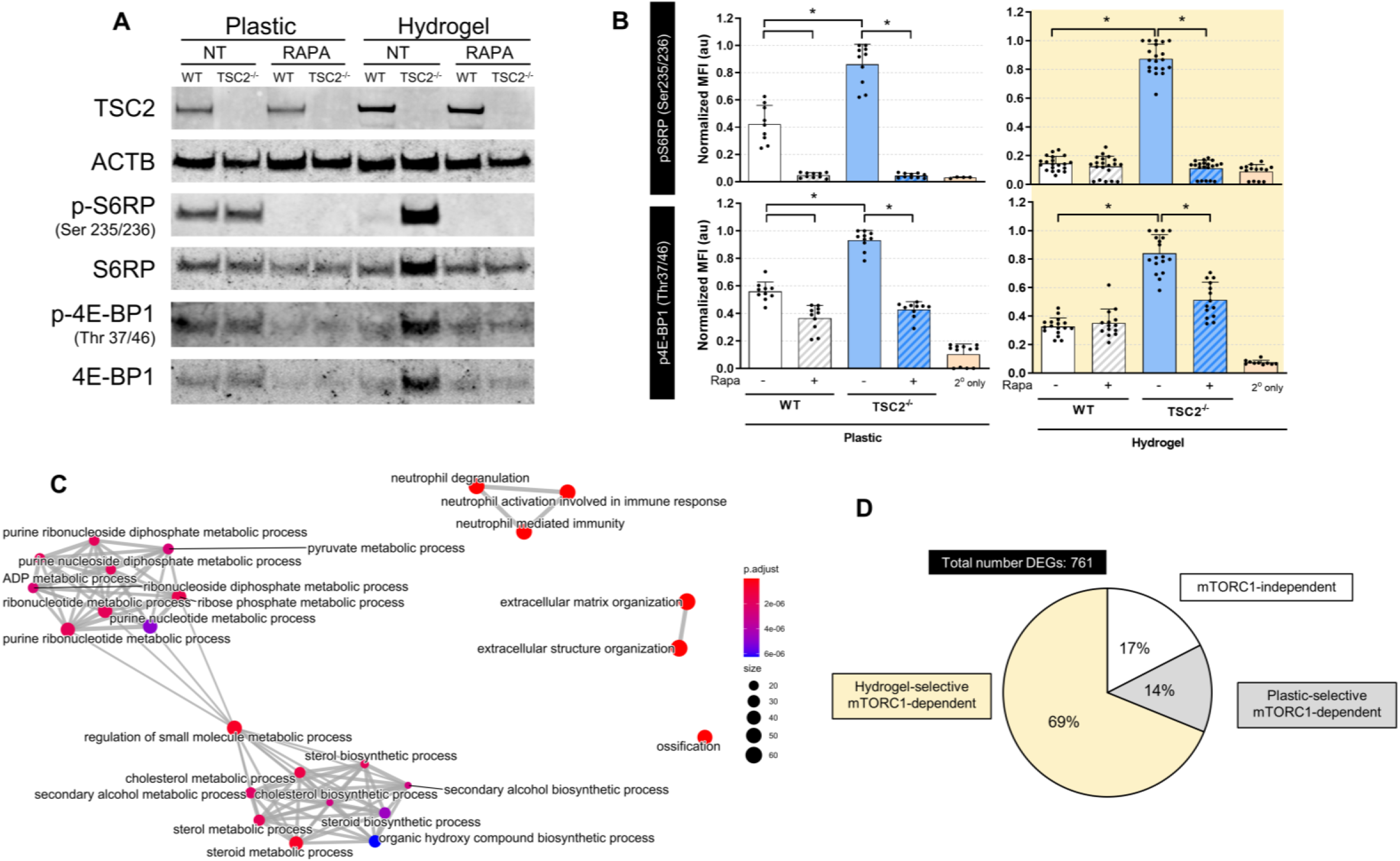
Hydrogel culture potentiates differential mTORC1-signalling between WT and TSC2^-/-^ cells. (**A**) Low input Western blot of protein collected from cells cultured for three days on plastic or hydrogel ± 20nM rapamycin. NT = no treatment, Rapa = rapamycin treatment. (**B**) Quantification of immunofluorescence values reported in normalized (scaled by replicate maximum value) mean fluorescence intensity. Each point indicates the mean fluorescence intensity from a well of cells cultured on hydrogel or plastic for three days ± 20nM rapamycin. Secondary only values are determined from wells probed with fluorescent secondary antibody only (mean ± SD; * = p < 0.05 by Student t-test; n = 9-20). (**C**) Network analysis of GO terms enriched in the list of DEGs found significant (FDR < 0.05) in the interaction between genotype and culture substrate. The 25 most significantly enriched terms are plotted. (**D**) Classification of DEGs according to pattern of expression across genotypes, ECM condition, and in the presence or absence of rapamycin. Gene clusters and classification scheme shown in Fig. S2B-C).

We sought to further explore the genotype-selective changes induced by hydrogel culture by interrogation of our bulk RNA-seq dataset. To do so, we tested for genes with a significant coefficient fit to the genotype:substrate interaction term (See Supplementary Materials and Methods) and identified 761 DEGs at FDR < 0.05 (Table S1C). Network analysis of GO terms enriched in this DEG list revealed two principal nodes, one related to sterol synthesis and the other to ribonucleotide metabolism (Fig. 2C, Table S2C). Notably, both these metabolic pathways have been associated with mTORC1 activity (*27*).

To unearth mTORC1-dependent transcriptomic alterations between WT and *TSC2^-/-^* that differ between plastic and hydrogel culture, we clustered the 761 DEGs based on their expression pattern across the 8 experimental conditions (Fig. S2B). Strikingly, genes related to sterol synthesis and ribonucleotide metabolism partitioned largely into two distinct clusters (Fig. S2B). We next classified each gene cluster into one of three categories based on the magnitude of expression differences between WT and *TSC2^-/-^*, and whether the expression changes were rescued by rapamycin (Fig. S2C). Remarkably, we find that 69% of the 761 DEGs showed a greater (or a unique) difference between WT and *TSC2^-/-^* cells in hydrogel culture compared to plastic, which was rescued by rapamycin (Fig. 2D). Together, these results demonstrate that hydrogel culture potentiates differential mTORC1 signalling between WT and *TSC2^-/-^* cells, reinforcing a physiologically relevant environment in which mTORC1-dependent phenotypes can be identified.

### Three-dimensional drug screen identifies compounds that modulate invasion and cell viability

We next employed our hydrogel culture system to identify potential therapeutic compounds. Cell death was measured at the single cell level by application of the live cell imaging fluorophore SyTOX, which selectively permeates cells with compromised plasma membrane integrity. We first tested a known cytotoxic compound, the proteasome inhibitor carfilzomib, and identified substantial cell death by live cell imaging (Fig. 3A, S3A). Additionally, we confirmed the ability to detect invasion modulation effects at the single cell level by employing the known anti-invasion Src kinase inhibitor dasatinib (Fig. 3B, S3B). To achieve the throughput necessary for a therapeutic screen, we acquired live cell images by high content microscopy paired with automated image analysis tools developed in house. We calculated the drug screen Z’ (a metric for assay quality) to be 0.873 for cytotoxicity measurements and 0.533 for invasion modulation. We subsequently screened a curated library of 800 structurally diverse, bioactive, membrane-penetrant compounds (Fig. 3C, Table S3A). Of these compounds, 39% have been trialed and shown to be safe for use in humans. We tested each compound on both WT and *TSC2^-/-^* cells in the presence and absence of rapamycin to elucidate mTORC1-dependency.

**Fig. 3.**
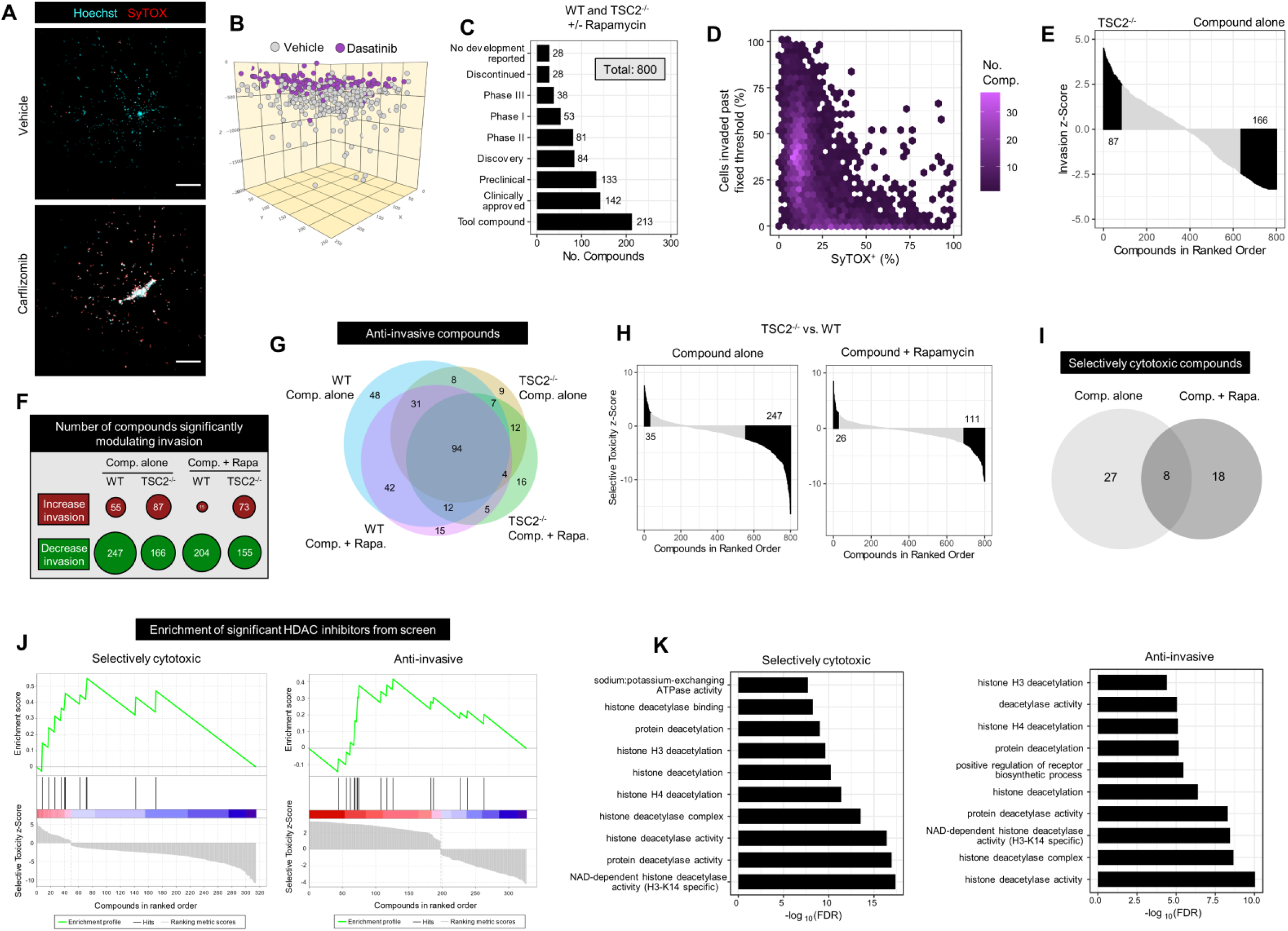
Three-dimensional drug screen identifies HDAC inhibitors as anti-invasive and selectively cytotoxic towards TSC2^-/-^ LAM cells. (**A**) Representative maximum intensity projection image of *TSC2^-/-^* cells in hydrogel culture for three days ± 200nM carfilzomib. Scale bars of 250µm. (**B**) Computational reconstruction of cellular spatial positions following three-day hydrogel culture of *TSC2^-/-^* cells ± 40nM dasatinib. Note that treated and untreated were in separate wells; cells were plotted in the same volume for ease of visualizing relative distances travelled. (**C**) Highest development status reported for the 800 compounds contained in the curated kinase inhibitor and tool compound libraries. A 3D drug screen was conducted on WT and *TSC2^-/-^* cells following three-day treatment with 5µM compounds ± 20nM rapamycin. (**D**) Compound invasion modulation plotted against cytotoxicity, aggregating results across genotype and rapamycin treatment. Fixed invasion threshold determined by median invasion distance of untreated controls. Hexagonal plot employed to demonstrate compound densities. (**E**) Waterfall plot of compound invasion z-scores in ranked order; positive values indicate invasion potentiation, while negative values indicate invasion attenuation. Compounds conferring statistically significant invasion modulation highlighted in black. Data presented for *TSC2^-/-^*, no rapamycin treatment condition. (**F**) Number of compounds significantly modulating invasion (potentiating or attenuating) for each genotype in the presence of absence of 20nM rapamycin. Bubble area proportional to number of statistically significant targets. (**G**) Overlap of compounds identified as anti-invasive in each listed condition. (**H**) Waterfall plots of compound selective toxicity z-scores in ranked order; positive values indicate increased cytotoxicity towards *TSC2^-/-^* cells, negative values indicate increased cytotoxicity towards WT cells. Compounds conferring statistically significant selective cytotoxicity highlighted in black. (**I**) Overlap of compounds identified to be selectivity cytotoxic towards *TSC2^-/-^* cells, with or without 20nM rapamycin. (**J**) Enrichment plot for compounds annotated to target HDACs, derived from an adapted implementation of GSEA. Hits (black vertical lines) in the red region indicate compounds with a favourable effect, hits in the blue region indicate compounds with an undesirable effect. (**K**) Top 10 most statistically significant GO terms. Analysis performed using targets identified as statistically significantly enriched in screen data by Elion^TM^ algorithm.

We found a wide variety of compounds with invasion modulatory and cytotoxic capabilities (Table S3B-C). Unsurprisingly, highly cytotoxic compounds also led to a reduction in bulk invasion (Fig. 3D). This trend was independent of genotype and rapamycin treatment (Fig. S3C). However, we observed many compounds which conferred an anti-invasive effect in the absence of detectable cytotoxicity (Fig. 3D, S3C). We next computed therapeutic invasion z-scores (i.e., statistical measure of compound effect size) by comparing against the vehicle control invasion distribution. Remarkably, while we identified several anti-invasive compounds, numerous compounds significantly increased invasion (Fig. 3E-F), a phenotype that would be otherwise overlooked if screening on two-dimensional plastic and could lead to severe adverse consequences in the clinical setting. In general, more compounds in this library were identified to significantly attenuate rather than potentiate invasion (Fig. 3F). Importantly, we observed a substantial overlap in the compounds identified to be anti-invasive across genotypes and treatment conditions, with very few drugs demonstrating a genotype-selective block to invasion (Fig. 3G, S3D). Together, these data demonstrate the identification of a collection of compounds which block invasion in these cell populations, irrespective of *TSC2* genotype.

A key goal in the therapeutic development landscape for LAM is the identification of compounds which exert selective cytotoxicity towards *TSC2^-/-^* cells. Interestingly, we observed that *TSC2^-/-^* cells exhibited pan-compound resistance, with over 7-fold more compounds demonstrating significant cytotoxicity towards WT compared to *TSC2^-/-^* cells (Fig. 3H). This selectivity is reduced to half with the addition of rapamycin, suggesting generalized resistance is largely due to mTORC1 hyperactivation in *TSC2^-/-^* cells. (Fig. 3H). We compared the list of compounds that are selectively cytotoxic towards *TSC2^-/-^* cells in the presence versus absence of rapamycin, and observed only a 15% overlap, indicating therapeutic vulnerabilities vary depending on mTORC1 signalling activity (Fig. 3I). In summary, we identified a suite of anti-invasive and selectively cytotoxic therapeutics which can be mined for further development in LAM (Table S3B-C).

### Enrichment analysis predicts HDAC inhibitors as anti-invasive and selectively cytotoxic towards *TSC2***^-/-^** cells

To refine our small molecule list for further investigation, we sought to identify outperforming compounds which modulate targets of the same class. Using the known annotated targets of the employed compounds, we performed target enrichment analysis by adapting the Gene Set Enrichment Analysis (GSEA) algorithm. We identified targets conferring well-established selective cytotoxicity towards *TSC2^-/-^* and anti-invasive classes, including proteasome inhibition (cytotoxicity) and Src and Rho kinase inhibition (anti-invasive) (Fig. S3E-F, Table S4A-D). Of note, Src inhibition, a therapeutic route explored in LAM, was found to be selectively cytotoxic towards WT cells (Fig. S3E). Remarkably, pan-HDAC inhibition was observed to be the only class in the top 10 most significant annotations for selective cytotoxicity towards *TSC2^-/-^* and generalized anti-invasion. We note a substantial favourable enrichment of HDAC-targeting compounds by both metrics, however, not all compounds annotated to inhibit HDACs performed favourably (Fig 3J).

A limiting factor to our analyses was the small number of compounds which were identified to selectively eliminate *TSC2^-/-^* cells. We sought to extend our compound list *in silico* using a structure-based approach with a mechanism of action prediction algorithm (Elion^TM^). In brief, chemical features are extracted from compound structures and matched with screen performance values to train a machine learning algorithm for prediction of other possibly efficacious compounds. Compounds predicted to be efficacious *in silico* are then analyzed by target enrichment and pathway analysis. Using this approach, we corroborated HDACs as highly enriched targets for both selective cytotoxicity and anti-invasion (Table S5A-B). GO term analysis on significant targets identifies nearly all top predicted pathways relate to deacetylation activity, for both selective cytotoxicity and anti-invasion (Fig. 3K, Table S5C-D). These data together highlighted HDAC inhibitors as promising therapeutics which we explored further and present herein.

### HDAC inhibitors are selectively cytotoxic towards *TSC2^-/-^* cells exclusively in hydrogel culture

We further tested 11 HDAC inhibitors from our compound library at a wider range of concentrations and identified three to be selectively cytotoxic towards *TSC2^-/-^* cells: SAHA (clinically, Vorinostat), SB939 (Pracinostat), and LBH589 (Panobinostat), all of which are pan-HDAC inhibitors (Fig. 4A). We note the atypical therapeutic dose-response curves and selectivity, demonstrating marginal differences in IC_50_ *per se* but substantial variation in maximal toxicity (Fig. 4B). Selective cytotoxicity was largely reversed by co-treatment with rapamycin, suggesting mTORC1-dependency. Importantly, the magnitude of cytotoxic selectivity between WT and *TSC2^-/-^* cells increased with treatment duration (Fig. S4A). We corroborated selective cell death functionally via clonogenic assays (Fig. S4B). Remarkably, when these HDAC inhibitors were tested with cells cultured on plastic, we did not observe any genotype-selectivity in their cytotoxic profile (Fig. 4A-B). In addition, inhibitor profiles employed in plastic culture did not change in the presence of rapamycin, suggesting a loss of mTORC1-dependency for cytotoxic effects (Fig. 4A-B). While HDAC inhibitors did modulate the proliferation of cells in hydrogel culture, a substantial proliferation blockade was exerted when cells were cultured on plastic, in both genotypes (Fig. S4C). Together, these data indicate a striking difference in cellular responses to HDAC inhibitor treatment while cultured on plastic versus hydrogel. Importantly, HDAC inhibitors only demonstrate mTORC1-dependent selective toxicity towards *TSC2^-/-^* cells while treated in hydrogel culture. These data are consistent with observation of hydrogel culture potentiating differential mTORC1 signalling between WT and *TSC2^-/-^* cells (Fig. 2, S2).

**Fig. 4.**
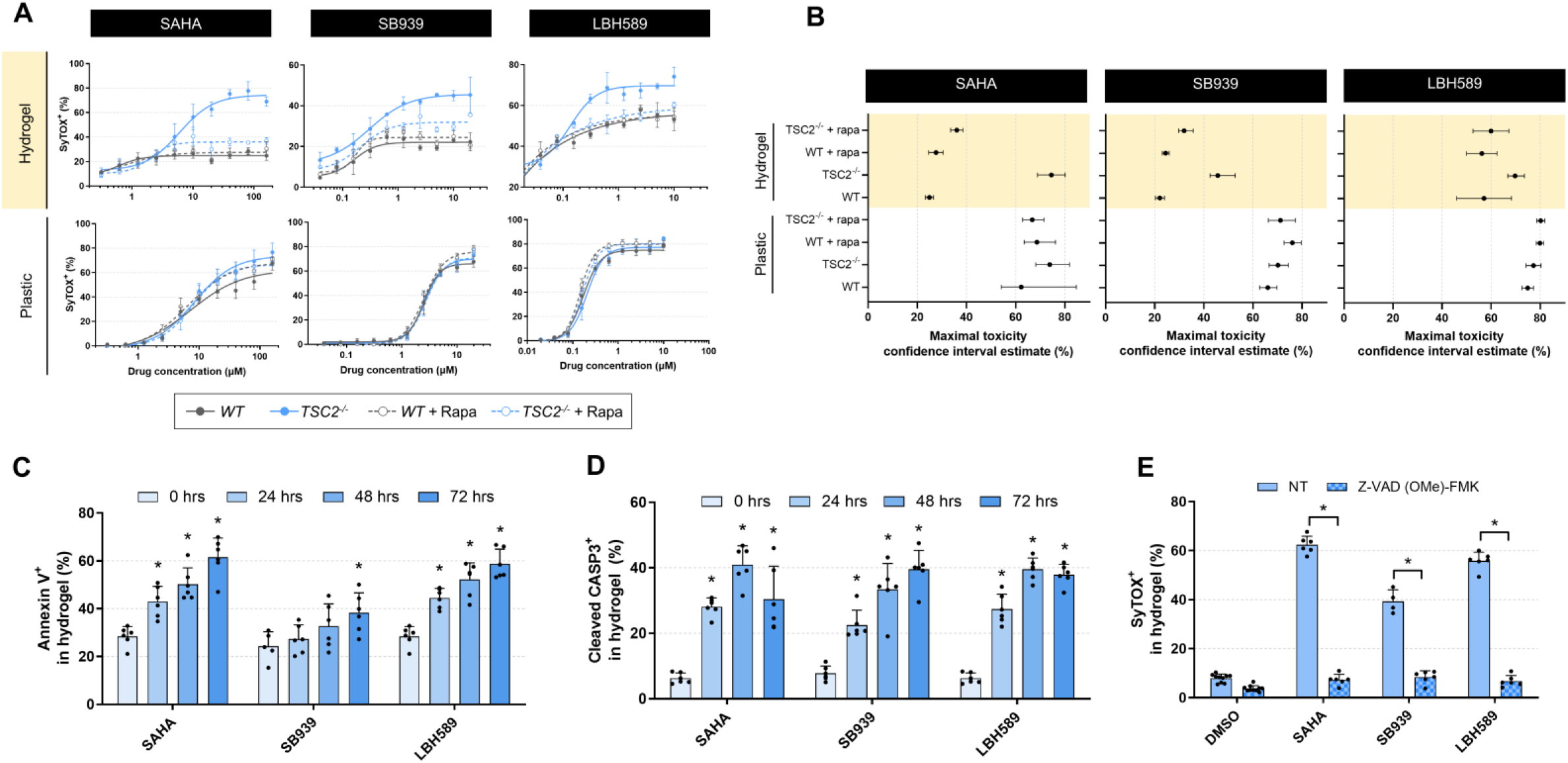
Three safe-in-human HDAC inhibitors induce mTORC1-dependent selective cytotoxicity exclusively in hydrogel culture. (**A**) Dose-response cytotoxicity curves of cells treated with the indicated HDAC inhibitor for three days while cultured on plastic or hydrogel ± 20nM rapamycin. Data fit via four-parameter logistic regression (mean ± SD; n = 3). (**B**) Confidence intervals of HDAC inhibitor maximal toxicity, estimated by four-parameter logistic regression models generated in A. (**C-D**) Quantification of *TSC2^-/-^* cells positive for (**C**) Annexin V or (**D**) cleaved caspase 3, following three-day treatment with HDAC inhibitors in hydrogel (20µM SAHA, 5µM SB939, and 1µM LHB589, mean ± SD; * = p < 0.05 by ANOVA with Dunnett post-hoc comparison to 0 hrs; n = 6). (**E**) Quantification of *TSC2^-/-^* cells positive for SyTOX following three-day HDAC inhibitor treatment (20µM SAHA, 5µM SB939, and 1µM LHB589) in hydrogel ± 25µM Z-VAD (OMe)-FMK (mean ± SD; * = p < 0.05 by Student t-test; n = 4-6).

### HDAC inhibitors induce cell death via apoptosis

We next sought to probe the mode of cell death induced by HDAC inhibitors. Previous studies have provided evidence for both HDAC inhibitor-mediated apoptosis as well as autophagic cell death (*28*). Considering we observed a reduction in cell death when co-treated with rapamycin, a potent inducer of autophagy, we hypothesized the predominant cell death mode to be apoptosis. To test this postulation, we employed live cell apoptosis imaging reagents, including cleaved caspase 3 (CASP3) and Annexin V. We validated their activity in our hydrogel culture using staurosporine, a known inducer of apoptosis (Fig. S4D-E). For all three HDAC inhibitors, we observed temporal accumulation of Annexin V and cleaved CASP3 with treatment duration in hydrogel (Fig. 4C-D, S4F). Importantly, we discerned a complete rescue of cell death by co-treatment with the caspase inhibitor Z-VAD (OMe)-FMK (Fig. 4E, S4G). Together, these data demonstrate the employed HDAC inhibitors induce apoptotic cell death in hydrogel culture.

### HDAC inhibitors are anti-invasive, independent of cytotoxic effects

To separate the anti-invasive effects from cytotoxic effects of these HDAC inhibitors, we identified and computationally removed SyTOX^+^ cells from invasion calculations (Fig. S5A). We determined all three HDAC inhibitors exhibited a dose-dependent anti-invasion effect on SyTOX^-^ cells (Fig. 5A-B). HDAC inhibitors exerted anti-invasive effects on both WT and *TSC2^-/-^* cells in the presence or absence of rapamycin. (Fig 5A-B, S5B-C). Of note, the effect size was generally larger in the *TSC2^-/-^* cells, and LBH589 demonstrated a trend towards reduced invasion that was not statistically significant. Remarkably, of the 11 HDAC inhibitors we tested, eight demonstrated anti-invasive effects in a dose-dependent manner (Fig. S5D). When aggregated as a class of therapeutics, there is a clear increase in anti-invasive effects with escalating doses, independent of cytotoxicity (Fig. 5C). Together, these data demonstrate HDAC inhibitors are effective anti-invasive agents independent of their cytotoxic profile.

**Fig. 5.**
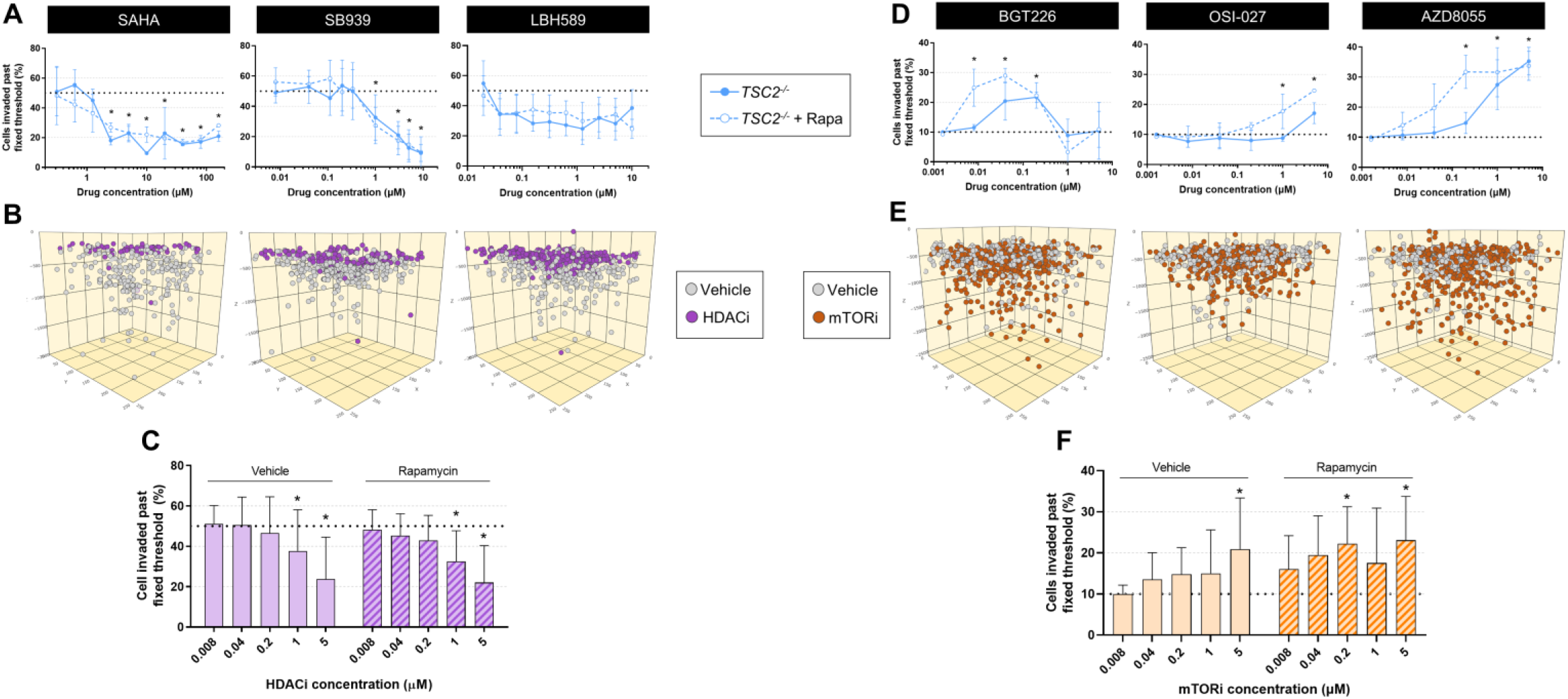
HDAC inhibitors attenuate cell invasion independent of cytotoxicity while mTOR inhibitors potentiate the invasion phenotype. (**A**) Live *TSC2^-/-^* cells invaded past fixed threshold set by median invasion distance of vehicle control, upon three-day HDAC inhibitor treatment ± 20nM rapamycin (mean ± SD; * = p < 0.05 by ANOVA with Dunnett post-hoc comparison to untreated; n = 3). (**B**) Computational reconstruction of live cell spatial positions upon three-day hydrogel culture of *TSC2^-/-^* cells ± HDAC inhibitor (HDACi) treatment (5µM SAHA, 5µM SB939, 1µM LBH589). Note that treated and untreated cells were in separate wells; cells were plotted in the same volume for ease of visualizing relative distances travelled. (C) Aggregated effect of 11 HDAC inhibitors on *TSC2^-/-^* live cell invasion ± 20nM rapamycin. Fixed threshold set by median invasion distance of vehicle control (mean ± SD; * = p < 0.05 by ANOVA with Dunnett post-hoc comparison to untreated; n = 33 via 11 HDACi, n = 3 each). (**D**) Live *TSC2^-/-^* cells invaded past fixed threshold set by 90^th^ percentile invasion distance of vehicle control, upon three-day mTOR inhibitor treatment ± 20nM rapamycin (mean ± SD; * = p < 0.05 by ANOVA with Dunnett post-hoc comparison to untreated; n =3). (**E**) Computational reconstruction of live cell spatial positions upon three-day hydrogel culture of *TSC2^-/-^* ± mTOR inhibitor treatment (40nM BGT226, 5µM OSI-027, 5µM AZD8055). (**F**) Aggregated effect of 3 mTOR inhibitors (mTORi) on *TSC2^-/-^* live cell invasion ± 20nM rapamycin. Fixed threshold set by 90^th^ percentile invasion distance of vehicle control (mean ± SD; * = p < 0.05 by ANOVA with Dunnett post-hoc comparison to untreated; n = 9 via 3 mTORi, n = 3 each).

### ATP-competitive mTORC1/2 inhibitors potentiate cell invasion

A surprising result of our 3D drug screen is the classification of ATP-competitive mTORC1/2 inhibitors as invasion potentiators (Fig. S3F). We interrogated this further due to its clinical relevance, as ATP-competitive inhibitors are in active development for a wide range of hyperactive mTOR conditions (*29*). Across a five-point dose-response curve, we observed an increase in invasion from three distinct mTORC1/2 inhibitors, independent of cytotoxic effects (Fig. 5D-E). Invasion potentiation was observed in both WT and *TSC2^-/-^* cells in the presence or absence of rapamycin (Fig. 5D-E, S5E-F). The extent of invasion potentiation varied across conditions: WT cells exhibited a greater increase in invasion compared to *TSC2^-/-^*, and the effect was exaggerated in both genotypes by co-treatment with rapamycin (Fig. S5G). Aggregating the effects of all three mTOR inhibitors showed a dose-dependent potentiation of invasion for this class of compounds (Fig. 5F). Together, these data demonstrate ATP-competitive inhibition of mTORC1/2 increases cell invasion.

### Xenotransplantation of LAM cell models into zebrafish larvae permits dynamic tracking of cell invasion

We next sought to evaluate the *in vivo* efficacy of the HDAC inhibitors SAHA, SB939, and LBH589. Consistent with previous findings, we found that loss of *TSC2* alone was insufficient to confer tumorigenicity upon subcutaneous xenotransplantation in immunodeficient mice (Fig. S6A). To avoid immortalization of our cell models-a process which dramatically alters cellular characteristics-we performed a well-established xenotransplantation assay in zebrafish larvae (*31, 32*). In this system, WT or *TSC2^-/-^* cells are injected into the hindbrain ventricle of zebrafish larvae 3 days post-fertilization, imaged 1 day post-injection (dpi) to ensure successful engraftment, and then imaged again at 4 dpi to visualize local invasion (Fig. 6A-B). Cells were tracked by their endogenous mCherry expression (*14*). The optical clarity of this system provides the advantage of enabling isogenic comparisons between WT and *TSC2^-/-^* human cells *in vivo* while dynamically tracking cell invasion.

**Fig. 6.**
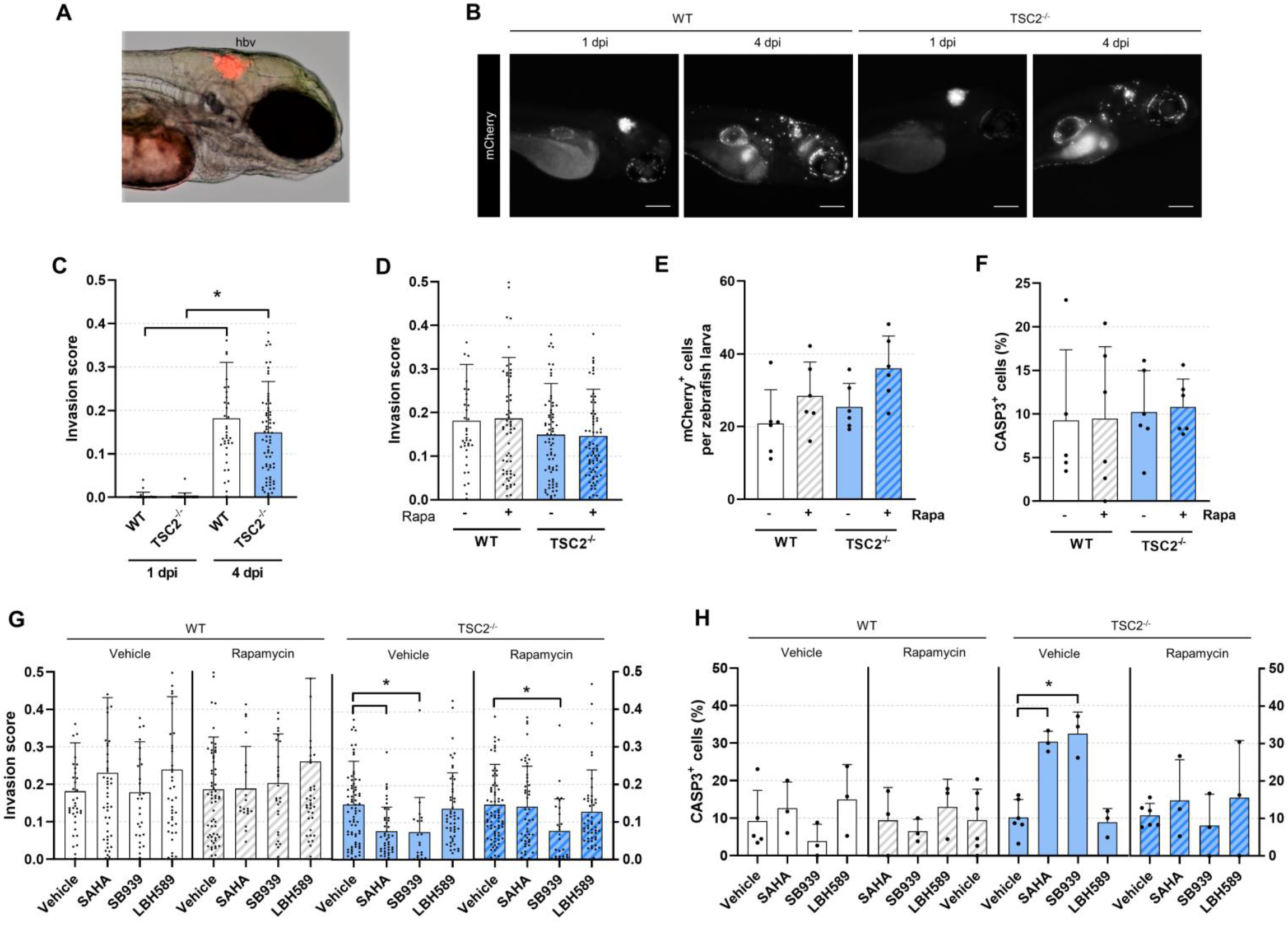
HDAC inhibitors are anti-invasive and selectively cytotoxic towards TSC2^-/-^ cells xenotransplanted into zebrafish. (**A**) Representative phase contrast image of 1 day post-injection (dpi) zebrafish larvae injected with *TSC2^-/-^* mCherry^+^ cells into the hindbrain ventricle (hbv). (**B**) Representative images of zebrafish larvae injected with mCherry^+^ WT or *TSC2^-/-^* cells into the hbv. Fish were imaged 1 and 4 dpi. Scale bars of 200µm. (**C-D**) Quantification of cell invasion using automated invasion analysis. Images analyzed in (D) were taken 4 dpi following three-day treatment ± 20nM rapamycin (mean ± SD; * = p < 0.05 by Mann–Whitney U test; n = 38 - 73). (**E**) Number of mCherry^+^ cells detected per zebrafish following whole larvae dissociation at 4 dpi and analysis by flow cytometry. Samples were treated for three days ± 20nM rapamycin. Each replicate is a pool of 15 – 20 zebrafish larvae (mean ± SD; * = p < 0.05 by Student t-test; n = 6). (**F**) Percentage of CASP3^+^ cells in the mCherry^+^ population from whole larvae dissociation at 4 dpi, following 3-day treatment ± 20nM rapamycin. Each replicate is a pool of 15 – 20 zebrafish larvae (mean ± SD; * = p < 0.05 by Student t-test; n = 5 – 6). (G-H) Effect of three-day HDAC inhibitor treatment (20µM SAHA, 5µM SB939, 1µM LBH589) ± 20nM rapamycin. (**G**) Invasion scores calculated on images acquired 4dpi (mean ± SD; * = p < 0.05 by the Kruskal-Wallis test with Dunn’s post-hoc comparison to vehicle treated; n = 27 – 73). (**H**) Percentage of CASP3^+^ cells in the mCherry^+^ population from whole larvae dissociation at 4 dpi. Each replicate is a pool of 15 – 20 zebrafish larvae (mean ± SD; * = p < 0.05 by ANOVA with Dunnett post-hoc comparison to vehicle treated; n = 3 – 6). Not all outliers in (C-D) and (G) are visualized due to trimmed axes (although outliers were included in mean ± SD and the statistical calculation).

To quantify invasion in an unbiased fashion, we computed the ratio of mCherry signal found outside the injection site compared to within (Fig. S6B-C). Using this method, we accurately detect near zero invasion scores 1 dpi, followed by a substantial increase 4 dpi (Fig. 6C). We observed comparable invasion scores between WT and *TSC2^-/-^* cells which were unaffected by rapamycin treatment, consistent with *in vitro* data (Fig. 6D). To quantify human cell proliferation and cell death, we digested and pooled whole larvae (15 - 20 per condition) followed by flow cytometry analysis, probing for mCherry and human-specific CASP3. Consistent with xenotransplantation in mice, these cells were not tumorigenic and the rate of clearance outstripped proliferation (Fig. S6D). The number of cells at 4 dpi was comparable between genotypes and unaffected by rapamycin treatment (Fig. 6E). The percentage of CASP3^+^ cells in the mCherry^+^ population was ∼10% and equivalent across conditions, similar to baseline cell death rates seen in hydrogel culture (Fig. 3A, 6F).

### HDAC inhibitors SAHA and SB939 block cell invasion and selectively eradicate *TSC2^-/-^* cells in vivo

We next employed our zebrafish xenograft system to assess the efficacy of HDAC inhibitors *in vivo*. To achieve the highest quality of pre-clinical evidence, experiments were conducted in a randomized, double-blinded, placebo-controlled fashion. We first established dose-toxicity profiles for each HDAC inhibitor: SB939 and LBH589 conferred an IC_50_ of 53.1 µM and 6.74 µM respectively, while the favourable toxicity profile of SAHA precluded calculation of an IC_50_ value (Fig. S6E). Of note, *in vivo* HDAC inhibitor potency correlated with the *in vitro* cytotoxicity profile. Zebrafish engrafted with either WT or *TSC2^-/-^* cells were treated with HDAC inhibitors by immersion therapy, in the presence or absence of rapamycin. Importantly, we used the same compound concentration as those employed *in vitro*, which was well below each compound’s IC_50_ value.

After three days of treatment, we observed that SAHA and SB939 exerted a statistically significant anti-invasive effect in the absence of rapamycin, exclusively on the *TSC2^-/-^* cells (Fig. 6G). SB939 also demonstrated a statistically significant anti-invasive effect in the presence of rapamycin. We note that live cells could not be distinguished from dead or dying cells in this quantification. However, by flow cytometry we observed an increase in the percentage of human *TSC2^-/-^* cells to be CASP3^+^ upon treatment with SAHA and SB939 (Fig. 6H) This effect was abrogated upon combination treatment with rapamycin and was not observed in the human WT cells. Together, these data indicate the HDAC inhibitors SAHA and SB939 exhibit *in vivo* anti-invasion and selective cytotoxicity effects towards *TSC2^-/-^* cells.

## DISCUSSION

Here, we subject novel tissue-engineered models of LAM to a three-dimensional drug screen to detect physiologically relevant therapeutics. We identified HDAC inhibitors as anti-invasive and selectively cytotoxic towards *TSC2^-/-^* cells, both *in vitro* and *in vivo*. In contrast, the gold standard therapeutic agent for LAM patients, rapamycin, did not exhibit any cytotoxic or anti-invasive effects. To our knowledge, this is the first high content compound screen to simultaneously track invasion modulation and cytotoxicity at the single cell level. Importantly, we report the first zebrafish xenotransplantation system using LAM-like cells, extending parametrization of therapeutic effects at the single cell level *in vivo* while comparing WT versus *TSC2^-/-^* cells.

Our research identifies pan-HDAC inhibitors as potential therapeutic candidates for pursuit in LAM patients. Notably, the selective cytotoxic effect towards *TSC2^-/-^* cells was only observed in hydrogel culture and would have otherwise been missed using standard screening in 2D tissue culture plastic (Fig. 4A-B). This finding complements a recent study evidencing therapeutic efficacy of HDAC inhibitors in a *Tsc1^-/-^*-driven mouse model of lymphangiosarcoma (*33*). HDAC inhibitors present an opportune class of molecules for pursuit due to the wide variety of compounds already approved for clinical use. Indeed, both SAHA and LBH589 are approved for use in cutaneous T cell lymphoma and multiple myeloma, respectively (*34, 35*). The safe-in-human toxicity profile of these compounds will facilitate rapid translation for testing in LAM patients. Importantly, our employed HDAC inhibitors exhibit selective cytotoxicity in an mTORC1-dependent manner, suggesting generalizable efficacy to mTORC1-driven malignancies. Of note, cutaneous T cell lymphoma cells have been observed to exhibit mTORC1 hyperactivation compared to matched normal controls (*36*).

In this work, we used equivalent HDAC inhibitor concentrations for *in vitro* and *in vivo* experiments; these concentrations were well-below dose-limiting toxicities in zebrafish (Fig. S6E). However, a critical outstanding question is whether the concentrations employed are physiologically attainable in humans. Pharmacokinetic studies of SAHA, SB939, and LBH589 in humans have demonstrated micromolar serum concentrations are achievable (*37–39*). In fact, the original pre-clinical work which formed the foundation for testing SAHA as a treatment in cutaneous T cell lymphoma used the drug *in vitro* at micromolar concentrations (*40*). Thus, we anticipate drug concentrations necessary to elicit a therapeutic effect are achievable in patients with LAM. We note that while LBH589 demonstrated a therapeutic effect *in vitro*, drug efficacy was not maintained upon *in vivo* testing, possibly due to altered bioavailability in the zebrafish. While SAHA and SB939 demonstrated *in vivo* efficacy, the majority of parameters assessed demonstrated mTORC1-dependency, similar to *in vitro*. Many LAM patients are on a chronic regime of rapamycin; thus, it is likely that clinical trials would require short-term withdrawal of rapamycin and acute treatment with HDAC inhibitors to elicit a therapeutic effect. Alternatively, these therapeutics may provide a benefit for LAM patients who are not currently treated with rapamycin, whether due to mild disease, intolerance, or resistance.

Our three-dimensional screening approach permitted the identification of many compounds with invasion modulatory capabilities. We encourage further mining of these data to uncover additional novel classes of therapeutics which modulate invasion and/or exert selectivity cytotoxic effects (Table S3B-C). As one example, we identified ATP-competitive inhibitors of mTOR to increase cellular invasion (Fig. 5D-F, S5E-G). This is of clinical significance and requires further investigation, considering this class of therapeutics is undergoing investigation in a broad range of oncogenic conditions (*29*).

Throughout our study, we note both genotype (loss of *TSC2*) and culture substrate (3D hydrogel) contribute to modelling features of LAM. Importantly, hydrogel culture potentiated differential mTORC1-signalling between WT and *TSC2^-/-^* cells, reinforcing a physiologically relevant environment in which mTORC1-dependent phenotypes can be identified (Fig 2, S2). However, we also note a variety of LAM features in our cellular models that exist independently from loss of *TSC2*. For example, cells isolated from both WT and *TSC2^-/-^* teratomas are equally invasive, present matching ACTA2^+^/PMEL^+^ profiles, and secrete similar levels of VEGF-D. Indeed, similar observations of LAM features in WT cells have been noted in a neural crest cell model (*14*). These data suggest perhaps, while loss of *TSC2* is critical for disease pathogenesis, the hallmark features of the putative “LAM cell” may already exist in a physiological, if not transient, context (e.g., during development, injury repair and inflammation). Critical consideration of the cell context is essential, even while employing isogenic comparisons, as different cell types exhibit distinct therapeutic vulnerabilities (*14*).

In summary, we have identified HDAC inhibitors as anti-invasive and selectively cytotoxic towards *TSC2^-/-^* cells *in vitro* and *in vivo*. While we have investigated three pan-HDAC inhibitors as potential candidates, our data points towards SAHA as the most efficacious against *TSC2^-/-^* cells while possessing the most favourable toxicity profile. On the path towards clinical translation, we anticipate testing of these compounds in diverse disease models. By validating compounds with orthogonal tools and techniques, we may elevate the most promising therapeutic for clinical trials.

## MATERIALS AND METHODS

### Study design

The objective of this research was to assess the LAM disease modelling capabilities of newly developed tissue-engineered cell models, and subsequently employ these models to identify novel therapeutic compounds. We conducted a 3D drug screen, and based on the acquired data, formulated and tested the following hypothesis: HDAC inhibitors are anti-invasive and selectively cytotoxic towards *TSC2^-/-^* cells. We employed a combination of *in vitro* and *in vivo* tools to test this hypothesis. Drug screen data was analyzed in a blinded, unbiased manner, and independently by two different researchers using distinct methods. Animal studies were conducted and analyzed in a double-blinded, randomized, placebo-controlled manner to generate the highest quality pre-clinical evidence. Blinding was achieved by codification of an investigator uninvolved in the experiments performed. A variety of experimental tools were employed to interrogate this hypothesis, described in subsequent Methods and in the Supplementary Materials and Methods. All reagents used and concentrations employed (if relevant) are reported in Supplementary Materials and Methods.

Sample sizes for both *in vitro* and *in vivo* studies were determined according to field-specific conventions. Power analysis was not employed. Data collection was not stopped prematurely, and every experimental replicate was analyzed. All data points were included in the data presentation; outliers were only excluded if there was definitive empirical evidence of technical error and noted as such in the figure legend. Experiments were repeated at least three times unless otherwise noted, with replicates collected at separate points in time and under independent conditions. RNA-seq data is accessible at the Gene Expression Omnibus (GEO) repository with accession GSE179044.

### Cell derivation and maintenance culture

LAM cell models were established via a previously reported *in vivo* differentiation protocol of human pluripotent stem cells (*23*). Briefly, we injected hPSCs into NSG mice to form teratomas, which were then explanted and expanded in smooth muscle-cell enriching conditions (Fig. S1A). We used a previously reported pair of mCherry^+^ WT and genome-engineered *TSC2*^-/-^ hPSCs for establishment of isogenic lines (*14*). Maintenance cultures were propagated on plastic containing a thin layer of Matrigel at 37°C, 5% CO_2_. hPSCs were cultured in Essential 8 medium and passaged in clumps by EDTA incubation, followed by cell scraping and wide-pore pipette transfer. LAM cells were cultured on Matrigel in Medium 231 containing Smooth Muscle Growth Supplement and passaged by 0.05% Trypsin as single cell suspensions.

### Hydrogel culture

Hydrogel culture was conducted according to a previously established protocol (*18*). Briefly, a hyaluronic acid polymer backbone was derivatized with methylfuran motifs (confirmed by ^1^H NMR) and conjugated to bismaleimide-terminated vitronectin and collagen-I-derived peptides, synthesized in house. Hydrogel viscoelasticity was increased by incorporation of methylcellulose derivatized with reactive thiol groups. Chemically synthesized hydrogel components were mixed and directly added to culture plates (384-well format) to gel at 37°C for 3 hours. Following gelation, wells were hydrated with PBS and then subjected to three media washes interspaced with incubations at 37°C for 45 mins. LAM cells were then dissociated, added to plates containing hydrogel, and centrifuged for 3 min. at 10*g* to achieve immediate contact with the hydrogel.

### Cell treatments and live cell staining

All compound treatments were conducted for 72 hours unless otherwise stated. See Supplementary Materials and Methods for 3D screen design, implementation, and analysis. Live cell imaging dyes (Hoechst, SyTOX, Annexin V, Cleaved Caspase 3) were incubated for 30 min. prior to imaging. Dyes were added as 10X concentrates in PBS; Annexin V diluent also contained 2.5mM CaCl_2_. To avoid cell detachment in the miniaturized well format, live imaging dyes were not washed prior to imaging; this did not impact image acquisition as dyes are minimally fluorescent unless bound to the target molecule.

### Microscopy

We employed a high content imager (Thermo Fisher Scientific, Arrayscan VTI) to acquire multi-well and multi-planar images. Whole-well images (384-well plate format) of cells invading through hydrogel, stained with live cell dyes, were acquired by widefield microscopy with 40µm interval z-stacks. Unstained cells were imaged using a brightfield module. Tiled images of cells grown and stained on plastic were also acquired by high content widefield microscopy. Rodent subcutaneous xenografts were visualized by *in vivo* imaging (PerkinElmer, IVIS®). Zebrafish larvae xenografts were imaged by epifluorescence widefield microscopy (Zeiss, AxioObserver 7). Image analysis methods are reported in Supplementary Materials and Methods.

### Animal studies

All animal experiments were conducted with approval from the University of Ottawa Animal Care Committee (Protocols #OHRI1666 and #CHEOe-3171), in accordance with the Canadian Council on Animal Care Standards and the Province of Ontario’s Animals for Research Act. NSG mice (Jackson Laboratory) were maintained in sterile housing conditions and fed autoclaved chow and water ad libitum. Adult *casper* (*41*) zebrafish (a gift from Dr. Leonard Zon, Boston Children’s Hospital, Boston, MA) were maintained in a recirculating commercial housing system (Aquatic Habitats, now Pentair) at 28°C in 14h:10h light:dark conditions in the aquatics facility at the University of Ottawa, Ottawa, ON. Adult *casper* zebrafish were bred according to standard protocol (*42*), and embryos were collected and grown in E3 medium (5mM NaCl, 0.17mM KCl, 0.33mM CaCl_2_, 0.33mM MgSO_4_) at 28°C in 10cm Petri dishes until the desired time point. Embryos were cleaned and provided with new media every 24hrs. See Supplementary Materials and Methods for additional experimental details.

### Statistical analysis

All figures are presented with individual data points (where graphically appropriate), with measures of central tendency and error to be mean and standard deviation, respectively, unless otherwise stated. Data pre-processing, statistical tests employed, sample number, and measured of central tendency and spread are reported in the figure legends. Two-sided tests were employed, and significance was attributed when p < 0.05. All analyses were of data from three independent experiments without removal of statistical outliers. Statistical tests employed were parametric except for analyses of zebrafish invasion data, where a non-normal distribution was observed. Calculation of drug screen statistics (e.g., z-scores, selectivity scores) are described in the Supplementary Materials and Methods.

## ACKNOWLEDGEMENTS

We thank Catherine Lawrence for her inspiration, members of our labs for continued insights, the Ottawa Bioinformatics Core Facility (particularly Christopher Porter) for assistance in processing RNA-seq data, and the Human Pluripotent Stem Cell Core Facility and Rima Al-awar (Ontario Institute for Cancer Research) for the custom designed drug library.

## Funding

Canadian Institutes for Health Research (CIHR) grant FRN-153188 (WLS) LAM Foundation pilot grant LAM0123P01-17 (WLS) Funds from Green Eggs and LAM (WLS and MSS) CIHR Vanier Canada Graduate Scholarship (AP) CIHR Tier 1 Canada Research Chair Program in Integrative Stem Cell Biology (WLS)

## Author contributions

Conceptualization: AP, JYL, SPD, RYT, JNB, MSS, WLS

Methodology: AP, JYL, NM, LJS, NA, CX, NM, LMJ, RYT, MSS, WLS

Investigation: AP, JYL, NM, NA, NM, EL, AC, CD Visualization: AP, JYL

Funding acquisition: MSS, WLS

Supervision: GM, LMJ, ASK, RYT, JNB, MSS, WLS

Writing – original draft: AP, WLS

Writing – review & editing: AP, JYL, NM, LJS, SPD, NA, CX, EL, AC, CD, GM, LMJ, ASK, RYT, JNB, MSS, WLS

## Competing interests

Authors declare that they have no competing interests.

## Data and materials availability

Processed data are available in the main text and the supplementary materials. Raw RNA-seq data is accessible at the GEO repository with accession GSE179044. Code written in house for data processing is available on request.

## SUPPLEMENTARY MATERIAL

### SUPPLEMENTARY MATERIALS AND METHODS

#### Study reagents and resources

Please see below (**Table 1**) for a list of key reagents and resources used in this study.

#### Cell culture

##### Pluripotent stem cell culture

H9 hPSCs (female) were maintained on a thin layer of 0.16 mg/mL Matrigel at 37°C, 10% CO_2_. Cells were fed with Essential 8 media, prepared in house. Cells were passaged by incubation with 500 µM EDTA for 3 min., then cell scraping and transfer to a new pre-coated plate by wide-bore pipette.

##### LAM and control cell model derivation and culture

LAM cell models were established via a previously reported *in vivo* differentiation protocol of human pluripotent stem cells (*23*). We differentiated a previously reported pair of mCherry^+^ WT and genome-engineered *TSC2*^-/-^ hPSCs, derived from the H9 parental lineage (female cells) (*14*). First, we generated teratomas in female NOD.Cg-*Prkdc^scid^ Il2rg^tm1Wjl^*/SzJ (NSG) mice as described in Mouse teratoma formation section. At end point, mice were euthanized and dissected under sterile conditions. The teratomas were extracted while carefully ensuring minimal mouse tissue remnants. The teratoma was minced and then rotated in a 5 U/mL Dispase solution at 37°C for 30 mins. Digested tissue was plated on a thin layer of 0.16 mg/mL Matrigel at 37°C, 5% CO_2_ in Medium 231 containing Smooth Muscle Growth Supplement. Tissue clumps were removed the following day. The remaining monolayer was expanded and passaged by treatment with 0.05% Trypsin for 5 min. Maintenance culture conditions included a thin layer of 0.16 mg/mL Matrigel at 37°C, 5% CO_2_ in Medium 231 containing Smooth Muscle Growth Supplement. Cells were expanded for two passages before cryopreservation and use in subsequent experiments at passages 3-5.

##### LAM and control cell model clonal isolation

LAM cells were clonally isolated by limiting dilution. Briefly, bulk cell cultures were dissociated and serially diluted to a concentration of ∼ 0.3 cells / 100 µL. We used this concentration to optimize number of single cells isolated while minimizing two or more cells contributing to a single clone. We added 100 µL of the suspension to each well of a 96-well plate containing a thin layer of 0.16 mg/mL Matrigel. Clones were expanded for 10 days before dissociating and plating onto the hydrogel.

#### Hydrogel culture

##### Reagent production

Hydrogel culture was conducted according to a previously established protocol (*18*). Briefly, a hyaluronic acid polymer backbone was derivatized with 5-methylfurfurylamine to 65% substitution (confirmed by ^1^H NMR). A vitronectin-mimetic peptide (maleimide)-KGGPQVTRGDVFTMPG, and MMP-degradable peptide crosslinker (maleimide)-KKGRGPQGIWGQKGPQGIWGQ-K(maleimide)S were synthesized using microwave-assisted Fmoc solid phase peptide synthesis with a CEM Liberty Blue automated peptide synthesizer. Hydrogel viscoelasticity was increased by incorporation of methylcellulose derivatized with reactive thiol groups.

**Table 1:**
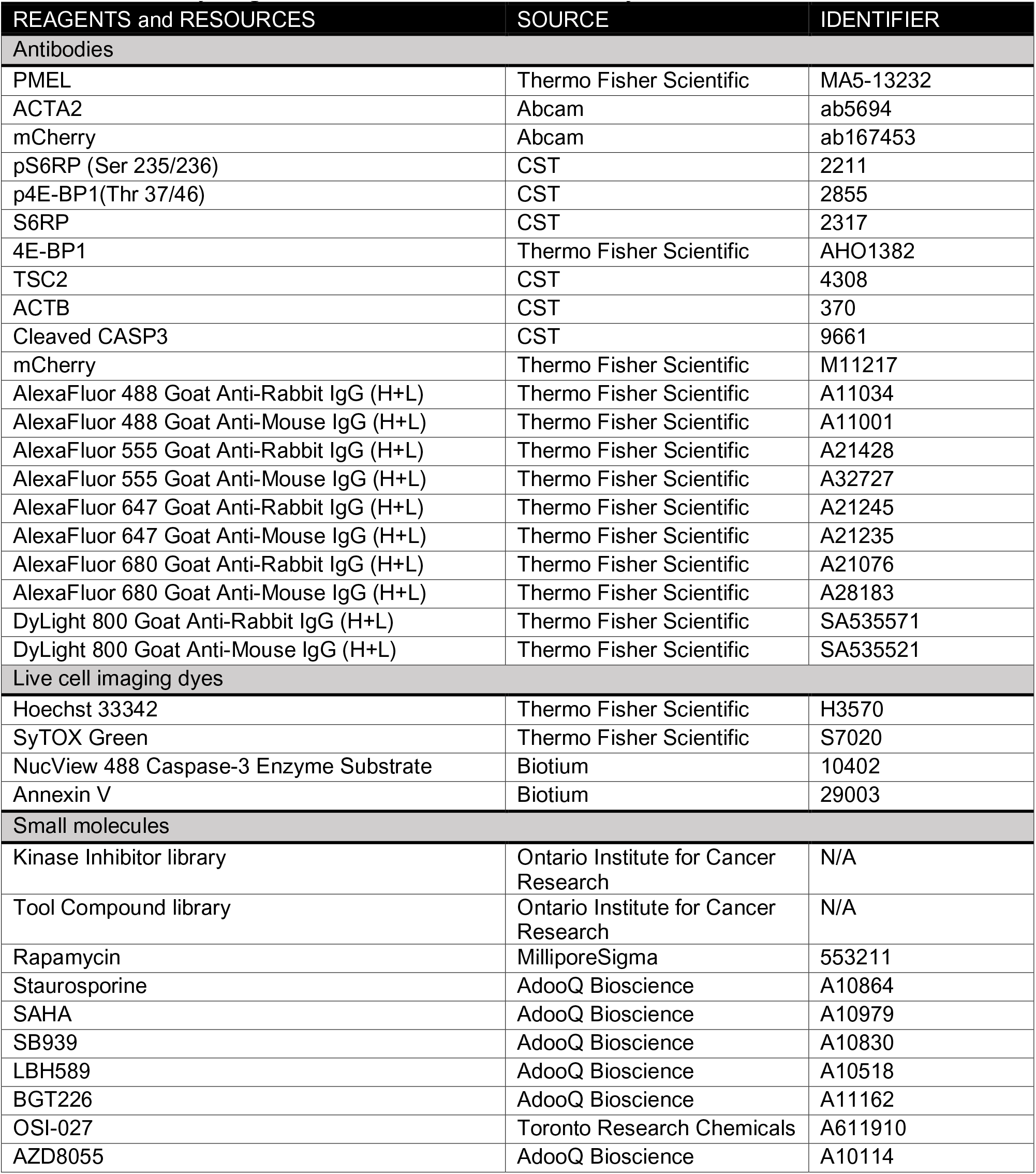
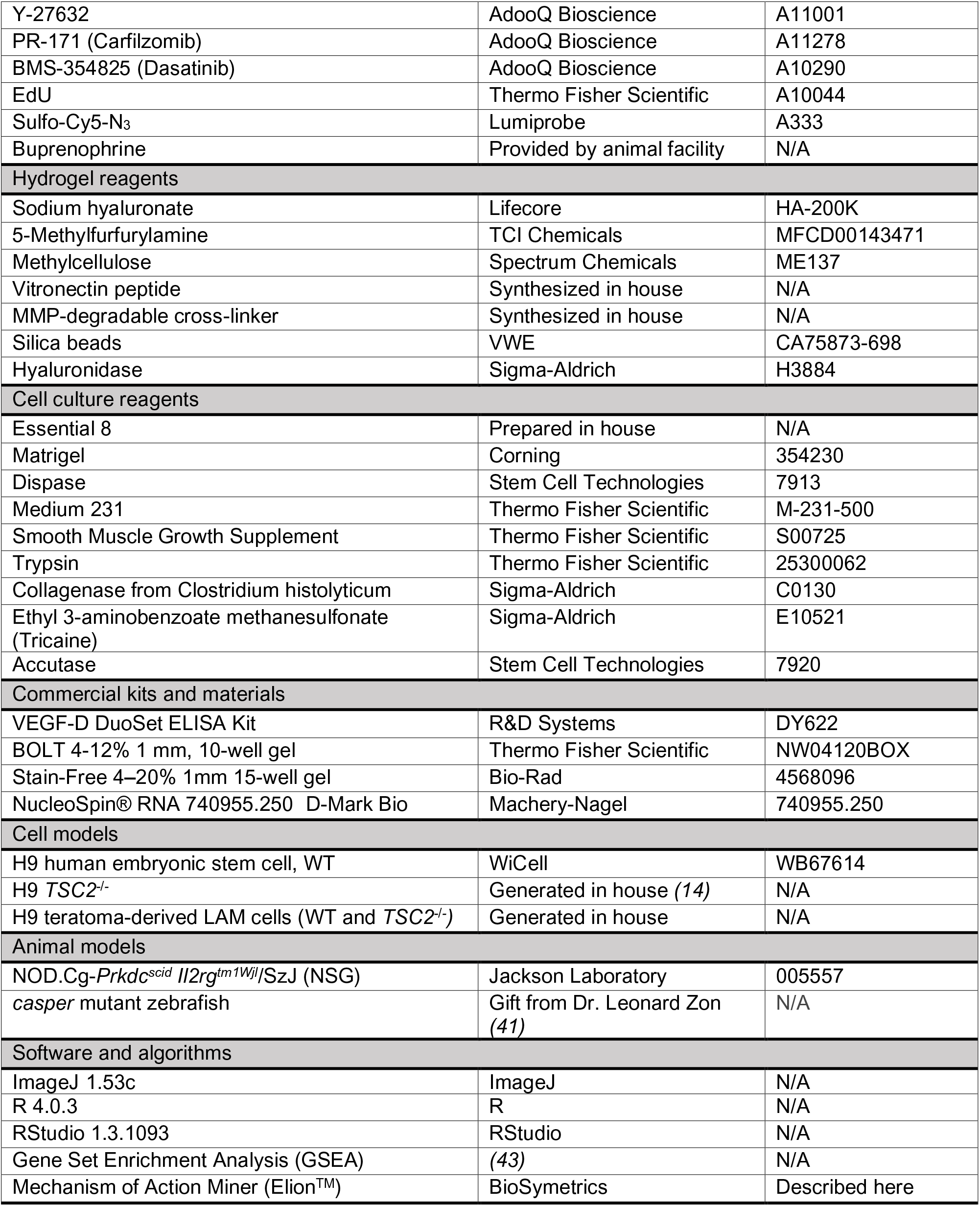
List of key reagents and resources used in this study.

| REAGENTS and RESOURCES | SOURCE | IDENTIFIER |
| --- | --- | --- |
| Antibodies |  |  |
| PMEL | Thermo Fisher Scientific | MA5-13232 |
| ACTA2 | Abcam | ab5694 |
| mCherry | Abcam | ab167453 |
| pS6RP (Ser 235/236) | CST | 2211 |
| p4E-BP1(Thr 37/46) | CST | 2855 |
| S6RP | CST | 2317 |
| 4E-BP1 | Thermo Fisher Scientific | AHO1382 |
| TSC2 | CST | 4308 |
| ACTB | CST | 370 |
| Cleaved CASP3 | CST | 9661 |
| mCherry | Thermo Fisher Scientific | M11217 |
| AlexaFluor 488 Goat Anti-Rabbit IgG (H+L) | Thermo Fisher Scientific | A11034 |
| AlexaFluor 488 Goat Anti-Mouse IgG (H+L) | Thermo Fisher Scientific | A11001 |
| AlexaFluor 555 Goat Anti-Rabbit IgG (H+L) | Thermo Fisher Scientific | A21428 |
| AlexaFluor 555 Goat Anti-Mouse IgG (H+L) | Thermo Fisher Scientific | A32727 |
| AlexaFluor 647 Goat Anti-Rabbit IgG (H+L) | Thermo Fisher Scientific | A21245 |
| AlexaFluor 647 Goat Anti-Mouse IgG (H+L) | Thermo Fisher Scientific | A21235 |
| AlexaFluor 680 Goat Anti-Rabbit IgG (H+L) | Thermo Fisher Scientific | A21076 |
| AlexaFluor 680 Goat Anti-Mouse IgG (H+L) | Thermo Fisher Scientific | A28183 |
| DyLight 800 Goat Anti-Rabbit IgG (H+L) | Thermo Fisher Scientific | SA535571 |
| DyLight 800 Goat Anti-Mouse IgG (H+L) | Thermo Fisher Scientific | SA535521 |
| Live cell imaging dyes |  |  |
| Hoechst 33342 | Thermo Fisher Scientific | H3570 |
| SyTOX Green | Thermo Fisher Scientific | S7020 |
| NucView 488 Caspase-3 Enzyme Substrate | Biotium | 10402 |
| Annexin V | Biotium | 29003 |
| Small molecules |  |  |
| Kinase Inhibitor library | Ontario Institute for Cancer Research | N/A |
| Tool Compound library | Ontario Institute for Cancer Research | N/A |
| Rapamycin | MilliporeSigma | 553211 |
| Staurosporine | AdooQ Bioscience | A10864 |
| SAHA | AdooQ Bioscience | A10979 |
| SB939 | AdooQ Bioscience | A10830 |
| LBH589 | AdooQ Bioscience | A10518 |
| BGT226 | AdooQ Bioscience | A11162 |
| OSI-027 | Toronto Research Chemicals | A611910 |
| AZD8055 | AdooQ Bioscience | A10114 |
| Y-27632 | AdooQ Bioscience | A11001 |
| PR-171 (Carfilzomib) | AdooQ Bioscience | A11278 |
| BMS-354825 (Dasatinib) | AdooQ Bioscience | A10290 |
| EdU | Thermo Fisher Scientific | A10044 |
| Sulfo-Cy5-N<sub>3</sub> | Lumiprobe | A333 |
| Buprenorphine | Provided by animal facility | N/A |
| **Hydrogel reagents** |  |  |
| Sodium hyaluronate | Lifecore | HA-200K |
| 5-Methylfurfurylamine | TCI Chemicals | MFCD00143471 |
| Methylcellulose | Spectrum Chemicals | ME137 |
| Vitronectin peptide | Synthesized in house | N/A |
| MMP-degradable cross-linker | Synthesized in house | N/A |
| Silica beads | VWE | CA75873-698 |
| Hyaluronidase | Sigma-Aldrich | H3884 |
| **Cell culture reagents** |  |  |
| Essential 8 | Prepared in house | N/A |
| Matrigel | Corning | 354230 |
| Dispase | Stem Cell Technologies | 7913 |
| Medium 231 | Thermo Fisher Scientific | M-231-500 |
| Smooth Muscle Growth Supplement | Thermo Fisher Scientific | S00725 |
| Trypsin | Thermo Fisher Scientific | 25300062 |
| Collagenase from Clostridium histolyticum | Sigma-Aldrich | C0130 |
| Ethyl 3-aminobenzoate methanesulfonate (Tricaine) | Sigma-Aldrich | E10521 |
| Accutase | Stem Cell Technologies | 7920 |
| **Commercial kits and materials** |  |  |
| VEGF-D DuoSet ELISA Kit | R&D Systems | DY622 |
| BOLT 4-12% 1 mm, 10-well gel | Thermo Fisher Scientific | NW04120BOX |
| Stain-Free 4-20% 1mm 15-well gel | Bio-Rad | 4568096 |
| NucleoSpin® RNA 740955.250 D-Mark Bio | Machery-Nagel | 740955.250 |
| **Cell models** |  |  |
| H9 human embryonic stem cell, WT | WiCell | WB67614 |
| H9 *TSC2*<sup>-/-</sup> | Generated in house (14) | N/A |
| H9 teratoma-derived LAM cells (WT and *TSC2*<sup>-/-</sup>) | Generated in house | N/A |
| **Animal models** |  |  |
| NOD.Cg-*Prkdc*<sup>scid</sup> *Il2rg*<sup>tm1Wjl</sup>/SzJ (NSG) | Jackson Laboratory | 005557 |
| *casper* mutant zebrafish | Gift from Dr. Leonard Zon (41) | N/A |
| **Software and algorithms** |  |  |
| ImageJ 1.53c | ImageJ | N/A |
| R 4.0.3 | R | N/A |
| RStudio 1.3.1093 | RStudio | N/A |
| Gene Set Enrichment Analysis (GSEA) | (43) | N/A |
| Mechanism of Action Miner (Elion™) | BioSymetrics | Described here |

##### Hydrogel gelation and culture

All chemically synthesized hydrogel components were mixed to the following final concentrations: 0.9 % methylfuronated hyaluronate, 2.3 mM MMP crosslinker, 100 µM vitronectin peptide, and 0.05 mg/mL thiolated methylcellulose. 15 µL of the solution was added to each well of a 384-well plate and permitted to gel at 37°C for 3 hours. Following gelation, wells were hydrated with PBS and then subjected to three media washes interspaced with incubations at 37°C for 45 mins. LAM or control cells were then dissociated, added to plates containing hydrogel, and spun for 3 min. at 10*g* to achieve immediate contact with the hydrogel.

#### Cell treatments

##### Drug treatments

All small molecule compounds were diluted in either DMSO or PBS, unless otherwise stated. The appropriate diluent-matched vehicle control was included in every experiment. Drug treatments were added directly to wells containing cells at 5X concentrations to avoid washing off any cells, particularly in sensitive miniaturized formats. Rapamycin was consistently used at a 20nM concentration. All compound treatments were conducted for 72 hours unless otherwise stated.

##### Live cell staining

Live cell staining dyes were used at the following final concentrations: 10 µg/mL Hoechst 33342, 50 nM SyTOX Green, 4 µM Caspase-3 Enzyme Substrate, and 0.2 µg/mL Annexin V. Dyes were incubated for 30 min. prior to imaging and added as 10X concentrates in PBS; Annexin V diluent also contained 2.5mM CaCl_2_. To avoid cell detachment in the miniaturized well format, live imaging dyes were not washed prior to imaging; this did not impact image acquisition as dyes are minimally fluorescent unless bound to the target molecule.

#### Cytotoxicity-invasion assay

Cells were permitted to invade through the hydrogel (384-well format) for 72 hrs while incubated at 37°C and 5% CO_2_. At end point, Hoechst and SyTOX were added directly to all wells as described in **Cell treatments** section. Whole-well multi-planar images were acquired by widefield microscopy with 40 µm separation between z-stacks. Following image acquisition, wells were fixed overnight in 10% formalin. We then added 1µg of silica beads to each well and acquired multiplanar brightfield images, with the plane of maximal contrast used to determine hydrogel-liquid interface (described in **Image analysis**). Acquiring location of the hydrogel interface (i.e., start of the cellular position) is essential for accurate invasion distance calculation; the hydrogel exhibits a meniscus which leads to a variable Z starting position depending on the XY location.

#### Three-dimensional drug screen

##### Screen design

Both WT and *TSC2^-/-^* cells were treated with every drug from the Ontario Institute for Cancer Research (OICR) Kinase Inhibitor and Tool Compound library (total of 800 compounds) at a concentration of 5 µM ± 20nM rapamycin. Cells were treated for 72 hr. while cultured in hydrogel and assessed at end point for cytotoxicity and invasion modulation as described in **Cytotoxicity-invasion assay**. Each plate included internal vehicle-treated only controls. Z’ was calculated for cytotoxicity and invasion modulation using vehicle-treated samples (negative control), 10 µM Y27632-treated (positive control, invasion), and 5 µM Carflizomib-treated (positive control, cytotoxicity).

##### Compound score calculation

To identify drugs with statistically significant effect(s), we computed z-scores for invasion modulation, cytotoxicity, selective invasion modulation, and selective cytotoxicity, for each WT and *TSC2^-/-^* in the presence or absence of rapamycin. We confirmed that the reference population of vehicle-treated controls for each metric was normally distributed and variance did not vary with effect mean. Cytotoxicity is determined by the percentage of SyTOX^+^ cells; invasion modulation is determined by the percentage of cells invading past the vehicle-control median threshold (See Image analysis section). Selective cytotoxicity is determined by the difference in cytotoxicity between WT and *TSC2^-/-^* cells, where positive values indicate more dead cells in the *TSC2^-/-^* condition. Selective invasion modulation is determined by the difference in cytotoxicity between *TSC2^-/-^* and WT cells, where positive values indicate fewer invading cells in the *TSC2^-/-^* condition. We then calculated p-values and corrected for multiple hypothesis testing by computing false discovery rates. All computation was performed using R 4.0.3 and RStudio 1.3.1093.

##### Target enrichment analysis

To refine our candidate compound list, we performed target enrichment analysis using a modified version of the GSEA algorithm (*43*). Enrichment analysis was performed separately for cytotoxicity and invasion modulation. For cytotoxicity, we focused our compound list to drugs that showed selective cytotoxicity, either in the presence or absence of rapamycin. If a drug was shown to be significantly beneficial in one condition but significantly detrimental in the other, it was excluded. We then derived a singular compound score by computing the arithmetic mean across the two conditions. Similarly, for invasion modulation, we focused our compound list to drugs that exhibit anti-invasion effects towards WT or *TSC2^-/-^* cells, either in the presence or absence of rapamycin. Again, we excluded compounds that showed opposing effects, and derived a singular compound score by arithmetic mean across conditions.

We next generated a background target list using known compound targets as annotated by the OICR. We created generalizable categories wherever possible, however, there were many targets that could not be grouped and conferred an n = 1 category. To avoid the possibility of bias, these categories were established by an independent author blinded to the original compound results. Using this background list and our compound score lists as described above, we determined target enrichment using the GSEA algorithm (*43*).

##### Elion^TM^ analysis

A limitation to our analyses is the small number of compounds which were identified to selectively eliminate *TSC2^-/-^* cells. We sought to extend our compound list *in silico* using a structure-based approach with Elion^TM^ (Mechanism of Action Miner), conducted by an independent group. Elion^TM^ is a software package that ingests binary phenotypic data linked to individual drug treatments to suggest possible underlying protein targets and molecular pathways. The platform inputs phenotypic screening data in the form of a two-column CSV file corresponding to the chemical structure in SMILES format alongside a binary bioactivity reading.

Using this dataset, a total of 8,000 features are generated for each supplied chemical structure. These features are comprised of chemical fingerprints (ECFP4, FCFP4, RDK-layered fingerprint, and MACCS) alongside physical properties (e.g., molecular weight, total polar surface area, LogP). If there are fewer than 8,000 rows in the input data set, a subset of features are chosen for downstream machine learning. The size of this feature subset is set to be 70% of the number of rows in the input data set. Feature selection is performed using a bootstrapped logistic regression strategy. In brief, a set number of features are sampled from the original feature set and are used to train a logistic regression model. The coefficients of this model are then used to rank feature importance. This process is repeated 5,000 times and the resulting coefficients are averaged for each feature to create a summarized feature importance score.

Once a feature set is chosen, a total of 6 machine learning models are built and evaluated on the input data set (XGBoost, random forest, Gaussian naive Bayes, uniform and distance weighted K-nearest neighbours, and Gaussian process classifiers). Each model is trained and evaluated using 10-fold cross validation while recording classification performance according to accuracy, ROC-AUC, precision and recall. The best performing model is then chosen and used to rank a set of 1 million compounds (curated from public databases) according to probability of inducing the given phenotype.

Of the 1M ranked compounds, several are annotated according to experimentally validated protein targets and mechanisms of action. The GSEA algorithm is used to determine which of these targets and MoAs are most positively enriched within the ranked set of compounds. We then subset this list using an FDR threshold to identify a set of enriched targets and MoAs. Using the enriched targets, we perform gene ontology and protein family pathway enrichment using the Fisher exact test. A Bonferroni corrected p-value threshold of 0.05 is used to identify cellular pathways corresponding to the phenotype of interest.

As a result of this process, Elion^TM^ translates an input phenotypic screen into three informative outputs. First, it supplies a ranked list of publicly available compounds prioritized according to their likelihood to induce the given phenotype. Second, it provides a list of targets and MoAs likely to mitigate the provided phenotype. Last, it annotates these targets with enriched cellular pathways. All of these results are presented in a web application annotated with rich descriptions and link-outs to relevant genetic databases.

#### Image analysis

##### Identification of cell spatial positions and cell invasion distance

We identified XYZ cell positions in the hydrogel by analysis of the Hoechst channel z-stack. We first determined XY positions by employing the ImageJ 1.53c “Find maxima” function on the z-stack maximum intensity projection. We automated the determination of the noise (or background) threshold by empirical iteration. Using the assumption that true Hoechst signal should be substantially above background fluorescence, we computed “Find maxima” with a liberal threshold, and then progressively increased threshold stringency until the number of identified points did not vary with each stepwise threshold change. Each maxima was determined to correspond to a single cell spatial location. Following, for each XY spatial position, we iterated through the Hoechst z-stack and identified to the point of maximal intensity, corresponding to the cell Z position.

To determine the invasion distance of each single cell, we must first know the cell starting position, which varied across XY positions due to the meniscus exhibited by the hydrogel. To identify determine hydrogel interface Z position across the XY plane, we used the silica bead brightfield images (described in Cytotoxicity-invasion assay). For each XY cell spatial position, we iterated through the brightfield z-stack and identified the point of maximal contrast, which corresponded to the layer containing silica beads (due to diffraction). We then determined individual cell distances travelled by computing the difference between cell starting and final positions. This process, automated for high throughput analysis, was scripted in ImageJ 1.53c.

##### Binarization of live cell stains

Live cell stains (i.e. SyTOX, Caspase-3 enzyme substrate, and Annexin V) were binarized into a positive or negative signal for each cell. We first created a masking around the Hoechst signal of each cell in the maximum intensity projection image, then measured the total fluorescent signal of the live cell stain within each masking. To binarize in an automated fashion, we fit an empirical probability density function (ePDF) by kernel density estimation on the vehicle control sample values. Assuming the majority of untreated samples should be negative for cell death stains, we determined the threshold for binarization to by the first local minimum of the negative control ePDF. Cells across conditions were then binarized according to their matched control threshold. This process, automated for high throughput analysis, was scripted in ImageJ 1.53c and R 4.0.3 within the RStudio 1.3.1093 environment.

##### Quantification of cellular invasion

Cellular invasion was determined by number of cells invading past a fixed distance. As the invasion distance varied slightly batch to batch (see Fig. 1F), distance thresholds for each experiment were based on within-experiment vehicle controls. We used both the median invasion of vehicle controls, which is sensitive to detecting decreases in invasion, and 90^th^ percentile invasion of vehicle controls, which is sensitive to detecting increases in invasion. Genotype-specific thresholds were employed due to the differing invasion distances between WT and *TSC2^-/-^* cells. To determine invasion of alive cells only, SyTOX^+^ positive cells were removed from the distribution prior to calculation of invasion percentages. This process, automated for high throughput analysis, was scripted in R 4.0.3 within the RStudio 1.3.1093 environment.

##### Immunofluorescence stain quantification

Immunofluorescence experiments were quantified using the raw image files, ensuring the absence of detector saturation. We created a masking around the Hoechst signal of each cell in the maximum intensity projection image, then measured the mean fluorescent signal of the protein-of-interest within the total masking area. Measurements per replicate were re-scaled from 0 – 1 by dividing by the replicate maximum value. We note that comparisons between plastic and hydrogel samples cannot be directly made, as the imaging parameters differ between the two-dimensional vs. three-dimensional environment (see Fig. 2B).

##### Invasion quantification upon zebrafish xenotransplantation

Invasion of transplanted mCherry^+^ cells was performed in a semi-automated fashion on blinded images (Fig. S6B). For each image, a region of interest was manually selected on the maximum intensity Z projection, to distinguish areas with mCherry^+^ cells from surrounding auto-fluorescent regions (e.g., zebrafish eye, yolk sac, ossicle). Images were then binarized using a constant threshold to distinguish positive signal from background. Pixels were classified into “invaded” or “not invaded” based on the distance from the center of the initial injection site. We used the average of the first local minima of the positive pixel histogram from 1 day post injected images to determine the distance for classification of invaded or not. Following pixel classification, the ratio of the total positive pixel intensity in each group was computed to determine the zebrafish invasion score. Groups were then unblinded and graphed. This process was scripted in ImageJ 1.53c and R 4.0.3 within the RStudio 1.3.1093 environment.

#### RNA-seq

##### RNA extraction and quality control

Extraction of RNA from cells embedded in hydrogel is made challenging by the low cellular density relative to the abundant extracellular matrix. To extract RNA, we developed an extraction protocol that combines phenol-chloroform phase separation with column-based purification. We first added TriZOL directly to wells and homogenized the cell-hydrogel mixture using a 26-gauge needle. We then centrifuged the lysate for 5min, 12,000*g*, at 4°C to pellet the cross-linked hyaluronic acid matrix. We extracted the supernatant and mixed in chloroform, followed by centrifugation to induce phase separation. The colourless aqueous phase was extracted and mixed with equal volumes of 70% EtOH. Following a brief incubation at room temperature, the solution was eluted through a Machery-Nagel Nucleospin column. We proceeded with column-based purification as per manufacturing protocol.

##### RNA-sequencing and raw data processing

RNA samples were shipped to the Donnelly Sequence Centre (Toronto, Canada) for RNA quality-control, library preparation, and next-generation sequencing. RNA integrity was assessed via Bioanalyzer (Agilent) and only samples with RIN > 8 were prepared for sequencing. Oligo(dT) priming via SMART-Seq v4 (Takara Bio) preparation kit was used to generate full-length cDNA libraries. Samples were subjected to paired-end sequencing on a NovaSeq 6000, 100c (Illumina) to a depth of ∼50 million reads per sample.

Raw sequence read quality was assessed by *FastQC*. Read feature assignments and duplication rates were determined using *featureCounts* and *Picard*. Overall mapping rate was assessed with *HISAT2*. Finally, read assignment to transcripts was performed using *Salmon*, generating a final pseudocount abundance matrix. All QC processing was summarized using *MultiQC* and programmed in R 4.0.3 within the RStudio 1.3.1093 environment. RNA-seq data is accessible at the Gene Expression Omnibus (GEO) repository with accession GSE179044.

##### Differential gene expression and enrichment analysis

Pseudocount abundance data generated by *Salmon* was imported into the *DEseq2* framework in R for differential expression and enrichment analysis. Principal components analysis was conducted on all samples to visualize transcriptomes in a two-dimensional space. For single variable differential expression testing, we subsetted samples to only include the untreated and fit the following model: *∼ batch + genotype + substrate*. We then tested for genes with significant coefficients by Wald test, separately for genotype and for culture substrate. To assess for changes across genotype that differs between matrix condition, we again subsetted for untreated samples and fit the following model: *∼ batch + genotype + substrate + genotype:substrate*. The interaction term coefficient for each gene was tested for significance by Wald test. Differentially expressed genes were called when false discovery rate (FDR) < 0.05, ± |log2FoldChange| > 1 (as indicated in the text).

To visualize expression values by heatmap or gene cluster, sample conditions were collapsed by abundance summation, normalized, and then transformed by regularized log_2_ transformation (implemented in *DEseq2*). Heatmaps were generated using the *pheatmap* package in R and gene clusters were generated by hierarchal clustering. GO term enrichment was performed using *clusterProfiler* on significant DEGs (FDR < 0.05, ± |log2FoldChange| > 1). All analysis was conducted using R 4.0.3 within the RStudio 1.3.1093 environment.

#### Animal studies

##### Mouse teratoma formation

hPSCs were dissociated into single cells by Accutase treatment for 15 min at 37°C. Single cells were harvested, washed, and resuspended in 5 mg/mL Matrigel. Female 8-week-old NSG mice were treated with buprenophrine 1 hour before injection, then anesthetized by isofluorane under a continuous stream of O_2_. We bilaterally injected 1×10^6^ hPSC into the mouse tibialis anterior. We allowed teratomas to grow over a 12-week period, after which mice were sacrificed and teratomas extracted.

##### Mouse subcutaneous xenografts

LAM cells were dissociated into single cells by 0.05% Trypsin treatment for 5 min. at 37°C, washed, and resuspended in 5 mg/mL Matrigel. Female 8-week-old NSG mice were anesthetized by isofluorane under a continuous stream of O_2_ and injected with 1×10^6^ cells subcutaneously in each rear flank. We monitored for palpable tumor growth weekly over a four-month period, after which animals were sacrificed.

##### Mouse IVIS image acquisition

Female 8-week-old NSG mice that were injected with LAM cells in each rear flank were monitored for tumor growth by endogenous mCherry expression of LAM cells. Mice were anesthetized by isofluorane under a continuous stream of O_2_ and shaved to eliminate background fluorescence from the fur coat. Mice were then imaged at fixed exposure times by *in vivo* imaging (PerkinElmer, IVIS®).

##### Zebrafish toxicity assay

72-hour post-fertilization (hpf) zebrafish larvae were arrayed one larva per well in a 96-well plate and treated with increasing concentrations of each inhibitor for 72 hrs to ascertain toxicity thresholds. There were no *in vivo* toxic effects at the experimental *in vitro* concentrations and thus, zebrafish experimental doses were chosen to stay consistent with *in vitro* treatment doses.

##### Zebrafish hindbrain ventricle xenotransplantation

For each injection experiment, a separate cryovial of cells was thawed and cultured 3 days prior to zebrafish transplantation, without any subculturing. On the day of transplantation, cells were dissociated by 0.25% trypsin, centrifuged for 5 mins at 300*g*, and resuspended in approximately 30 µL of culture medium for injection. 72 hpf zebrafish larvae were anesthetized with 0.09 mg/mL tricaine (Millipore Sigma) and arrayed in troughs of an agarose injection plate and used for cell transplantation using protocols described previously (*44, 45*). The cells were backloaded into a pulled capillary needle and allowed to settle for approximately 20 mins at 35°C to ensure a cell pellet at the bottom of the needle. A PLI-100A Pico-liter Microinjector (Warner Instruments) was used to manually inject 50-100 cells into the hindbrain ventricle (HBV) of each larva. Following injections, the larvae were kept at 35°C for the remainder of the experiment.

##### Zebrafish drug treatments

1 day post injection (dpi), injected larvae were screened on an Axio Observer 7 fluorescent microscope under an mCherry filter to ensure the presence of cells only in the HBV. Groups of 20-30 positively injected larvae were randomized into groups to be treated with either vehicle control (DMSO), 20 nM rapamycin alone, 5 µM SB939 alone, 20 µM SAHA alone, 1 µM LBH589 alone or with one HDACi in combination with rapamycin by immersion therapy for 72hrs. At the experimental endpoint (3 days post-treatment) the groups of larvae were blinded and imaged on the Axio Observer 7 using the z-stack function to capture cell movement in all planes. Blinded groups of images were then subjected to automated invasion analysis by an independent study author.

##### Zebrafish whole larval dissociation and fixation

At 1 dpi (baseline) and 4 dpi (three days post-treatment), 20 larvae from each group were euthanized and dissociated in 100mg/mL collagenase solution for approximately 30 mins. Upon completion of dissociation (i.e., single cell suspension formed), 200 µL of 100% FBS was added to slow the enzymatic reaction. The samples were then centrifuged for 5 min. at 300 *g* and the supernatant was removed, leaving a pellet of human tumor cells among the zebrafish cells. The samples were washed once in 30% FBS in PBS and centrifuged once more for 5 min. at 300 *g*. The supernatant was removed and 250µL of 4% PFA in PBS was added to each sample for 20 min. in the dark. 1mL of PBS was added and samples were centrifuged and PFA supernatant was removed. Samples were resuspended in 500 µL PBS, stored at 4°C, and blinded prior to flow cytometry analysis.

#### Immunofluorescence staining

The following protocol is for immunofluorescence staining of cells in monolayer culture on plastic. Modifications for whole-mount (WM) staining of cells in three-dimensional hydrogel are indicated throughout.

Cells were fixed with 4% PFA for 15 min. (WM: 30 min.) at room temperature. Wells were washed 3 x 5 min. (WM: 20 min.) with PBS, then permeabilized with 0.1% Triton-X in PBS for 20 min. (WM: 40 min) at room temperature. Wells were washed 3 x 5 min. (WM: 20 min.) with PBS, then blocked with 1% BSA in PBS for 1 hr. (WM: 2 hr.) at room temperature. We then added primary antibody diluted in blocking solution for overnight incubation at 4°C. The following concentrations of antibodies were employed: PMEL (1:50), ACAT2 (1:100), pS6RP^Ser235/236^(1:100), and p4E-BP1^Thr37/46^(1:200). The next day, wells were wash 3 x 5 min. (WM: 5 x 30 min.) with PBS, then incubated with fluorescent secondary antibodies diluted blocking solution for 1 hr. (WM: 2 hr.) at room temperature. All fluorescent secondary antibodies were used at a 1:1000 dilution. Wells were then washed 3 x 5 min. (WM: 5 x 30 min.) with PBS then counterstained with 10 µg/mL Hoechst 33342 for 30 min. (WM: 45 min.) Wells were washed 3 x 5 min (WM: 5 x 30 min.), then mounted with a 90% glycerol (WM: PBS, as the hydrogel disintegrates in glycerol) solution made in house, prior to imaging.

#### Enzyme-linked immunosorbent assay (ELISA)

Maintenance cultures of cells at equivalent densities were incubated for 16 hr. in Medium 231 ± 20nM rapamycin, without serum supplement. Conditioned media was collected and centrifuged to remove any cellular debris, then assayed by VEGF-D ELISA kit (R&D Systems, DY622) following the manufacturer protocol.

#### Flow cytometry

LAM cells were dissociated into single cells by 0.05% Trypsin treatment for 5 min. at 37°C, washed, and then fixed with 4% PFA for 15 min. at room temperature. Fixing solution was diluted out 1/10 in PBS, cells were pelleted by centrifugation, and supernatant discarded. For details on zebrafish single cell preparation, see Zebrafish whole organism dissociation and fixation section. Fixed single cell suspensions were permeabilized with 0.1% Triton-X in PBS for 20 min. at room temperature. Permeabilizing solution was diluted out 1/10 in PBS, cells were pelleted by centrifugation, and supernatant discarded. Samples were then blocked with 1% BSA or 5% Goat Serum in PBS for 1 hr. Cells were pelleted by centrifugation, supernatant discarded, and primary antibodies diluted in blocking solution were added for overnight incubation at 4°C. The following concentrations of antibodies were employed: PMEL (1:50), ACAT2 (1:100), mCherry (1:1000), and Cleaved CASP3 (1:500). The next day, primary antibody solution was diluted out 1/10 in PBS, cells were pelleted by centrifugation, and supernatant discarded. Samples were then incubated with fluorescent secondary antibodies diluted blocking solution for 1 hr. at room temperature. All fluorescent secondary antibodies were used as a 1:1000 dilution. Secondary antibody solution was diluted out 1/10 in PBS, cells were pelleted by centrifugation, and supernatant discarded. Samples were next counterstained with 10 µg/mL Hoechst 33342 for 20 min. at room temperature. Hoechst 33342 solution was diluted out 1/10 in PBS, cells were pelleted by centrifugation, and supernatant discarded. Finally, cells were strained and analyzed using the LSRFortessa (BD) flow cytometer.

#### Low input western blot

Hydrogel culture must be performed in a miniaturized format to maintain the appropriate mechanics as previously reported (*18*). Naturally, this poses a challenge for collecting sufficient protein for standard molecular biology methods, such as a western blot. To address this challenge, we developed a method for a low input western blot that includes in-well lysis and sample preparation, followed by a gel-based method for sample normalization.

Samples were cultured on plastic or in hydrogel for 72 hr. Following, sample media was aspirated to the hydrogel interface (leaving a similar volume in plastic wells) and an equivalent volume of 2X Laemmli-RiPA buffer was added to each well. Samples were incubated for 10 min. at 37°C and triturated up and down, careful not to disturb the hydrogel. The sample volume was then extracted and boiled for 10 min. at 70°C. As this extraction contains a large amount of non-cellular derived protein components (due to degradation of the hydrogel MMP-cleavable crosslinkers), standard protein quantification by colorimetric methods (e.g., BCA, and Bradford) are not reliable. Instead, we performed total protein quantification on gel-separated samples. First, an aliquot of each sample was electrophoresed on a stain-free 4–20% 1 mm 15-well gel, along with a serial dilution of a sample of known concentration. The gel was then activated by UV exposure and total protein visualized by ChemiDoc Gel Imager (Bio-Rad). We then calculated individual sample concentrations by comparing against the within-gel standard curve, without including bands corresponded to the hydrogel MMP peptides.

After sample extraction and quantification, we analyzed samples following standard western blotting procedures. To maximize sample input, Thermo Fisher Scientific BOLT gels were used, which contain space for up to 60 µL of sample per lane. We first separated samples by SDS-PAGE using a BOLT 4-12% 1 mm 10-well gel and MES running buffer. Samples were transferred onto a 0.45 µm PVDF membrane overnight at 4°C by wet transfer with Towbin buffer (containing 0.025% SDS and 10% MeOH). The membrane was then blocked by 5% BSA in PBS-T (0.1% Tween-20) for 1 hr. at room temperature. Following, the membrane was incubated overnight at 4°C in primary antibodies diluted in blocking buffer at the following concentrations: pS6RP^Ser235/236^(1:5000), p4E-BP1^Thr37/46^(1:1000), S6RP (1:500), 4E-BP1 (1:1000), TSC2 (1:5000), and ACTB (1:5000). The membrane was washed 3 x 5 min. with PBS-T, and then incubated for 1 hr. at room temperature in fluorescent secondary antibodies diluted 1:10,000 in blocking buffer. The membrane was washed 3 x 5 min. with PBS-T and then imaged using Odyssey Gel Imager (LI-COR Biosciences).

#### EdU proliferation assay

Cells were pulsed with 5 µM of EdU for 3 hr. Subsequently, cells were fixed with 4% PFA for 15 min. at room temperature. Wells were washed 3 x 5 min. with PBS, permeabilized with 0.1% Triton-X in PBS for 20 min. at room temperature, then washed again 3 x 5 min. We prepared the click reaction by mixing the following components in the described order, in PBS, to the indicated final concentrations: 4 mM Cu_2_SO_4_, 5 µM Sulfo-Cy5-N_3_, and 100 mM L-ascorbic acid. The click reaction mix was added to wells containing cells and incubated at room temperature for 30 min. Wells were washed 3 x 5 min. with PBS, counterstained with 10 µg/mL Hoechst 33342 for 30 min, washed again 3 x 5 min., and then imaged.

#### Clonogenic assay

Cells were plated on the hydrogel and treated with HDAC inhibitors at the designated concentration for 72 hr. Following treatment, wells were washed 3 x 20 min. with media to remove the drug from solution. The hydrogel was then solubilized by addition of 150 U hyaluronidase per 15 µL hydrogel and incubated for 1 hr. at 37°C. Following, 0.05% Trypsin was added to the wells for 10 min. at 37°C to dissociate cells. Wells were triturated and then plated on two-dimensional tissue cultures plates, in serial dilution. Cells were permitted to proliferate for 10 days, forming colonies from single cells. Following, wells were fixed with 4% PFA for 15 min. at room temperature, then washed 3 x 5 min. with PBS. Colonies were stained with 0.1% crystal violet for 1 hr. at room temperature, washed 3 x 5 min. with ddH_2_O, air dried, and imaged.

### SUPPLEMENTARY FIGURES

**Fig. S1.**
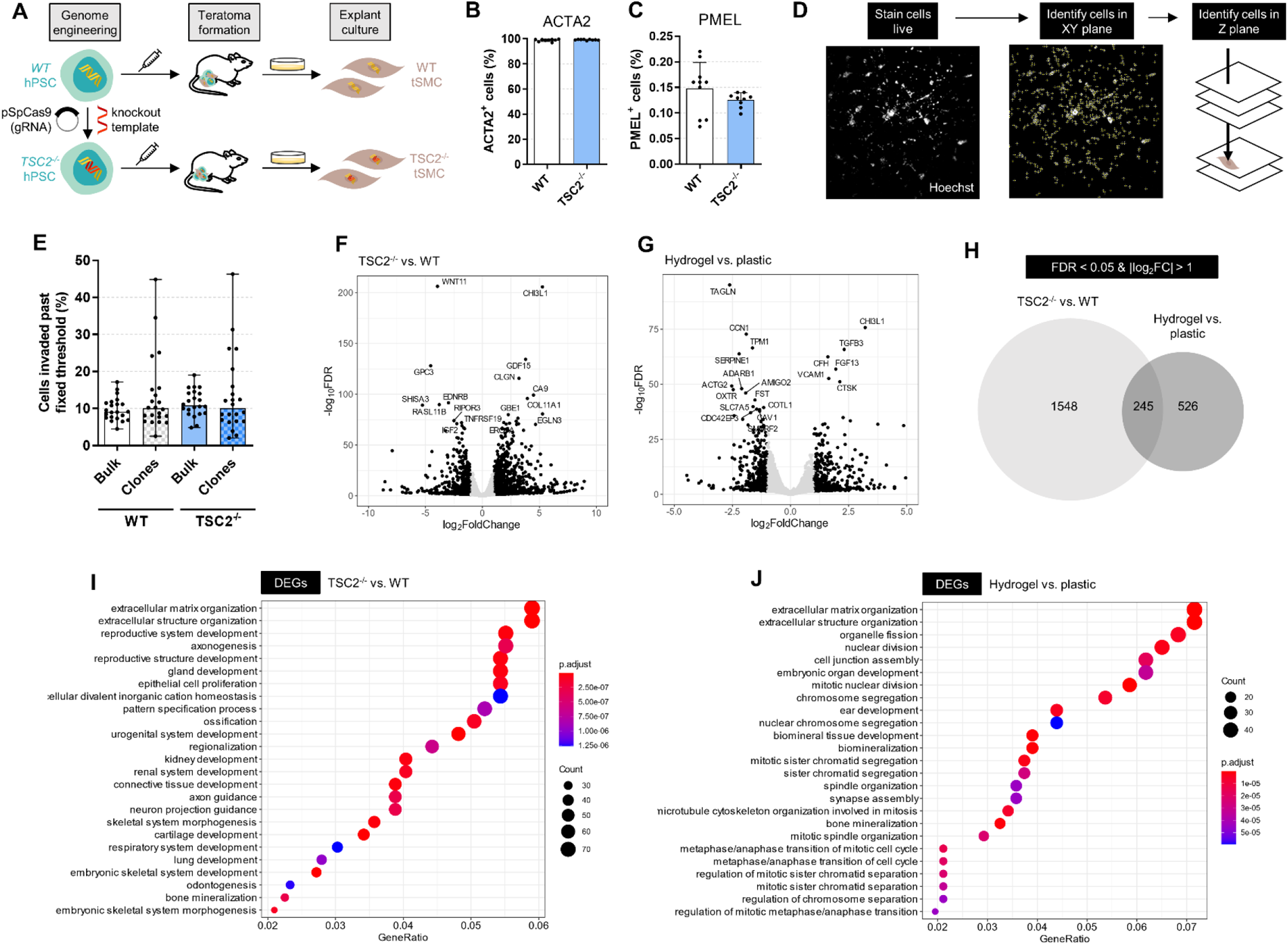
Hydrogel culture of stem cell-derived disease models exhibits features of LAM. (A) Schematic of generation of LAM cellular models. (B-C) Quantification of LAM markers by flow cytometry from cells in maintenance culture (mean ± SD; * = p < 0.05 by Student t-test; n = 10). (D)) Schematic of cell position identification in XYZ planes. (E) Percentage of cells invaded past threshold set by 90^th^ percentile invasion distance of bulk cultures, following three-day hydrogel culture. Bulk cultures are maintenance cultures of LAM cell lines; clones are populations of cells expanded from a single cell isolated from maintenance cultures prior to seeding on hydrogel (mean ± data range; no statistical test). (F-G) Volcano plot upon comparing *TSC2^-/-^* vs. WT cells (F) and hydrogel vs. plastic samples (G). Points highlighted in black are considered differentially expressed (FDR < 0.05 and |log_2_FC| > 1). The 20 most significantly DEGs are noted. (H) Overlap in DEG between genotype and culture substrate gene lists; genes considered as DEGs if FDR < 0.05 and |log_2_FC| > 1. (I-J) Dotplot of GO term enrichment analysis of DEG lists (FDR < 0.05 and |log_2_FC| > 1) upon comparing *TSC2^-/-^* vs. WT cells (I) and hydrogel vs. plastic samples (J). The 25 most significantly enriched terms are plotted.

**Fig. S2.**
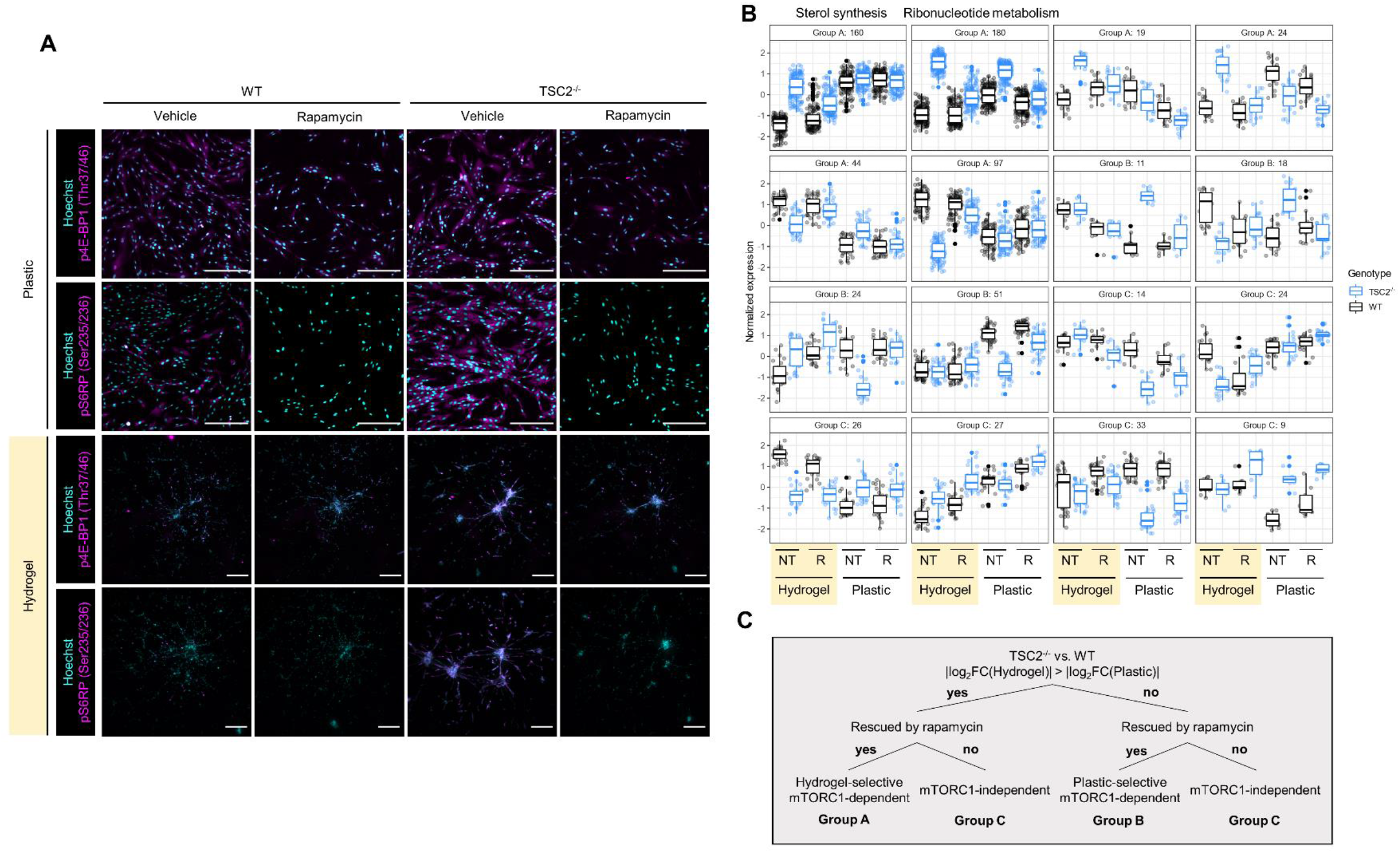
Hydrogel culture potentiates differential mTORC1-signalling between WT and TSC2^-/-^ cells. (A) Representative maximum intensity projection images used for quantification, following culture on hydrogel or plastic for three days ± 20nM rapamycin. Scale bars of 250µm. (B) Gene clusters following hierarchal clustering of DEGs found significant (FDR < 0.05) in the interaction between genotype and ECM (761 genes). Clustering was based on the pattern of gene expression across the 8 employed conditions. The first two clusters are annotated to be enriched in sterol synthesis and ribonucleotide metabolism terms. (C) Classification scheme applied to the gene clusters visualized in (B). Labelling of Groups A, B, and C is for ease of visualization in (B) and does not confer any specific biological meaning.

**Fig. S3.**
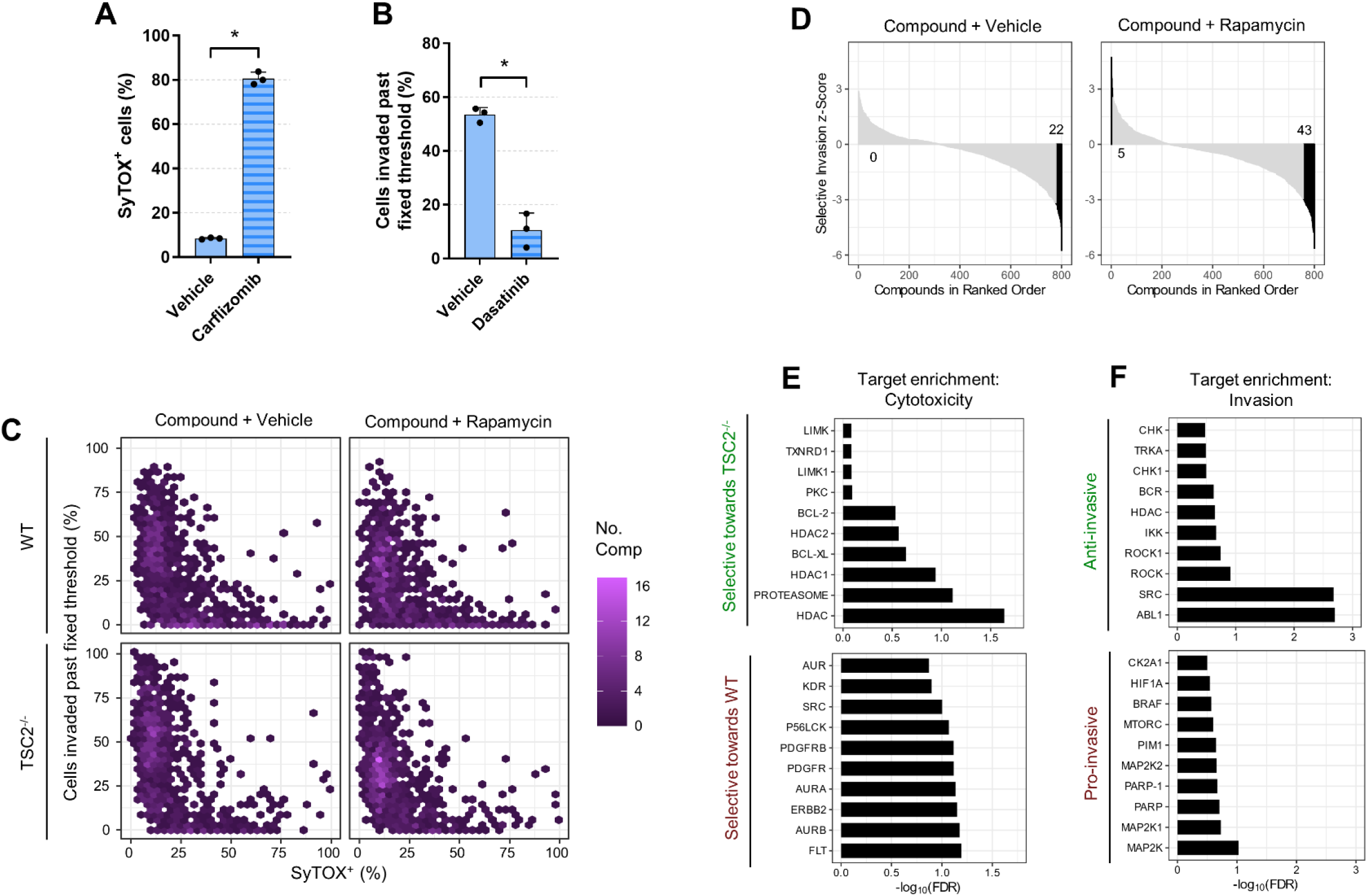
Three-dimensional drug screen identifies HDAC inhibitors as anti-invasive and selectively cytotoxic towards TSC2^-/-^ LAM cells. (A) Percentage of SyTOX^+^ *TSC2^-/-^* cells in hydrogel culture for three days ± 200nM carfilzomib (mean ± SD; * = p < 0.05 by Student t-test; n = 3). (B) Percentage of *TSC2^-/-^* cells invaded past fixed threshold (determined by median invasion distance of untreated controls), following three-day hydrogel culture ± 40nM dasatinib (mean ± SD; * = p < 0.05 by Student t-test; n = 3). (C) Compound invasion modulation plotted against cytotoxicity, separated by genotype and rapamycin treatment. Fixed threshold determined by median invasion distance of genotype-specific untreated controls. Hexagonal plot employed to demonstrate compound densities. (D) Waterfall plots of compound selective invasion z-scores in ranked order; positive values indicate greater anti-invasive effects towards *TSC2^-/-^*, negative values indicate greater anti-invasive effects towards WT. Compounds conferring statistically significant selective invasion modulation highlighted in black. (E-F) Top 10 most statistically significant targets enriched in screen data, stratified by screen parameter. Enrichment analysis was performed via adaptation of the GSEA algorithm, using annotated targets of the compound library.

**Fig. S4.**
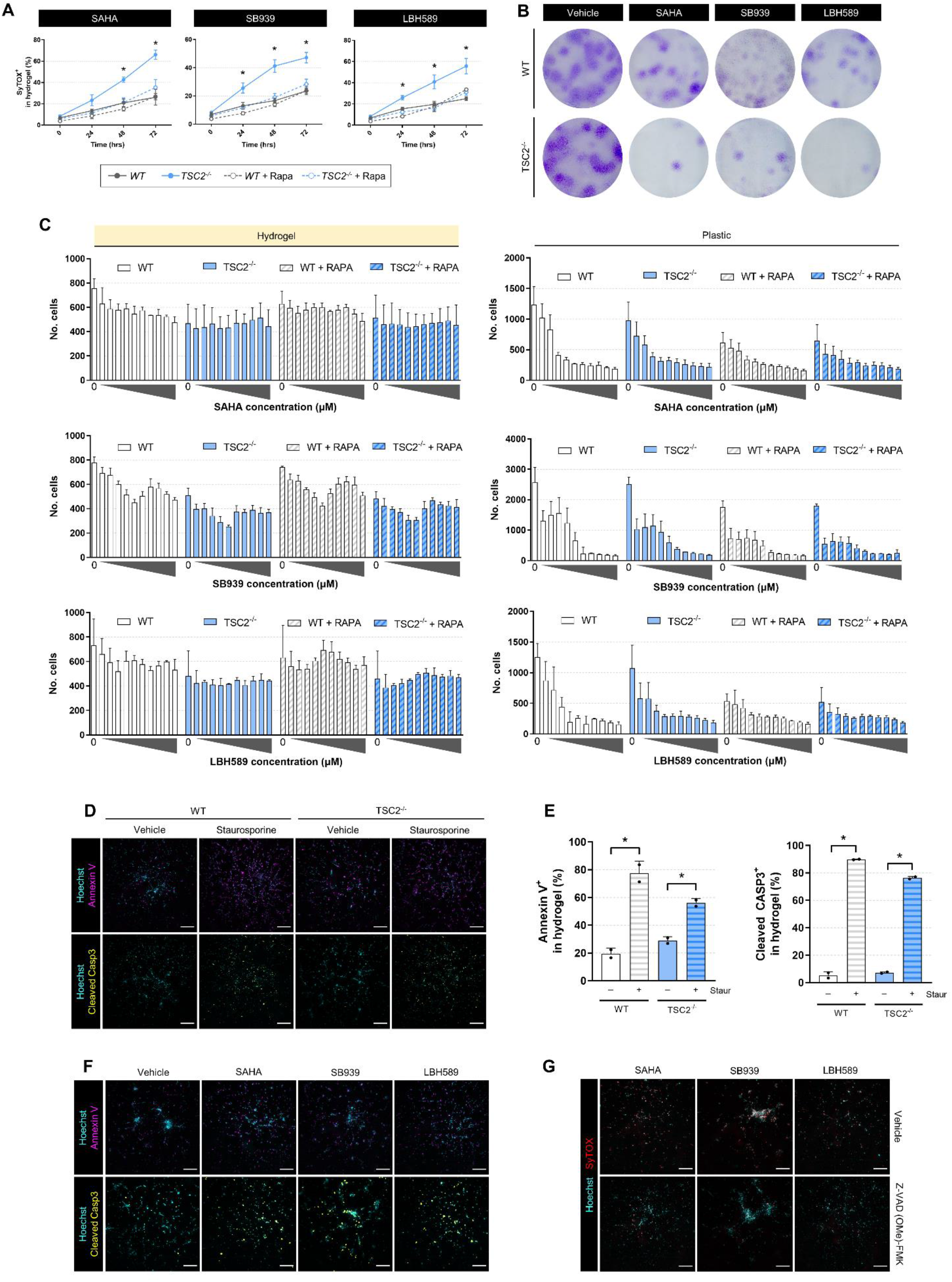
Three safe-in-human HDAC inhibitors induce mTORC1-dependent selective cytotoxicity exclusively in hydrogel culture. (A) Percentage of SyTOX^+^ cells following time-lapse HDAC inhibitor treatment (20µM SAHA, 5µM SB939, 1µM LBH589) of cells cultured in hydrogel ± 20nM rapamycin (mean ± SD; * = p < 0.05 by two-factor ANOVA with Tukey’s post-hoc comparison; n = 3). (B) Clonogenic assay following three-day HDAC inhibitor treatment (20µM SAHA, 5µM SB939, 1µM LBH589) of cells cultured in hydrogel. After treatment, cells were extracted from hydrogel and replated in 2D to assess clonogenicity. (C) Number of cells detected in culture by high content imaging following three-day HDAC inhibitor treatment in hydrogel or plastic culture ± 20nM rapamycin. Inhibitor concentrations escalated in two-fold increments: SAHA (0.31µM min, 160µM max), SB939 (0.04µM min, 20µM max), and LBH589 (0.02µM min, 10µM max). Mean ± SD, n = 3. (D-E) Representative maximum intensity projection images and quantification of live cell imaging dyes used in hydrogel culture, following 4hr treatment of 1µM staurosporine (mean ± SD; * = p < 0.05 by Student t-test; n = 2). Scale bars of 250µm. (F-G) Representative maximum intensity projection images of live cell imaging dyes used in hydrogel culture, following three-day treatment with HDAC inhibitors (20µM SAHA, 5µM SB939, 1µM LBH589) ± 25µM Z-VAD (OMe)-FMK. Scale bars of 250µm.

**Fig. S5.**
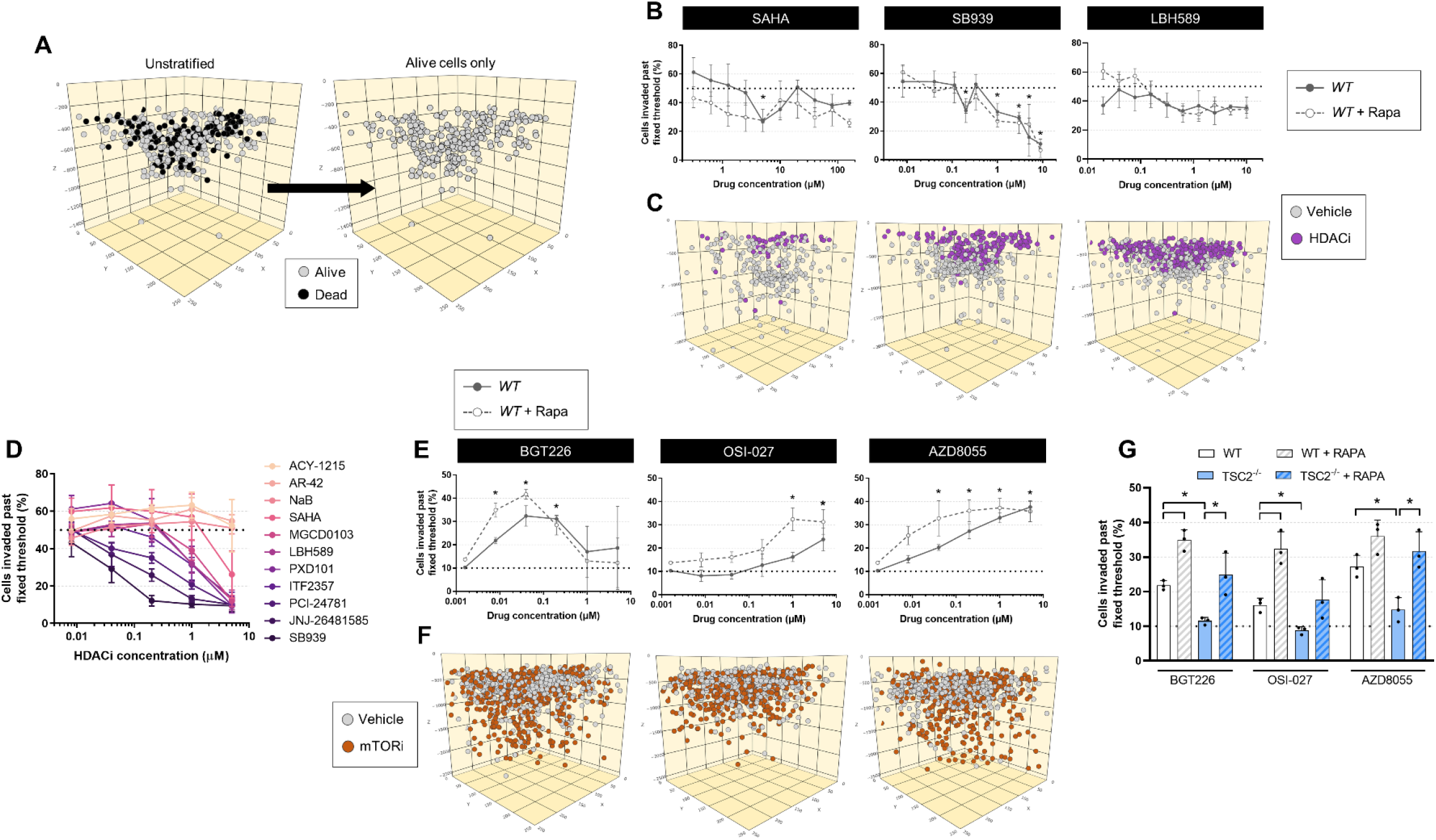
HDAC inhibitors attenuate cell invasion independent of cytotoxicity while mTOR inhibitors potentiate the invasion phenotype. (A) Schematic for removal of SyTOX^+^ cells to determine invasion distribution of live cells. (B) Live WT cells invaded past fixed threshold set by median invasion distance of vehicle control, upon three-day HDAC inhibitor treatment ± 20nM rapamycin (mean ± SD; * = p < 0.05 by ANOVA with Dunnett post-hoc comparison to untreated; n = 3). (C) Computational reconstruction of live cell spatial positions upon three-day hydrogel culture of WT ± HDAC inhibitor treatment (5µM SAHA, 5µM SB939, 1µM LBH589). Note that treated and untreated were in separate wells; cells were plotted in the same volume for ease of visualizing relative distances travelled. (D) Effect of 11 HDAC inhibitors on *TSC2^-/-^* live cell invasion ± 20nM rapamycin. Fixed threshold set by median invasion distance of vehicle control. (E) Live WT cells invaded past fixed threshold set by 90^th^ percentile invasion distance of vehicle control, upon three-day mTOR inhibitor treatment ± 20nM rapamycin (mean ± SD; * = p < 0.05 by ANOVA with Dunnett post-hoc comparison to untreated; n = 3). (F) Computational reconstruction of live cell spatial positions upon three-day hydrogel culture of WT ± mTOR inhibitor treatment (40nM BGT226, 5µM OSI-027, 5µM AZD8055). (G) Live cells invaded past fixed threshold set by 90^th^ percentile invasion distance of genotype-matched vehicle control, upon three-day mTOR inhibitor treatment (8nM BGT226, 1µM OSI-027, 200nM AZD8055) ± 20nM rapamycin (mean ± SD; * = p < 0.05 by Student t-test; n = 3).

**Fig. S6.**
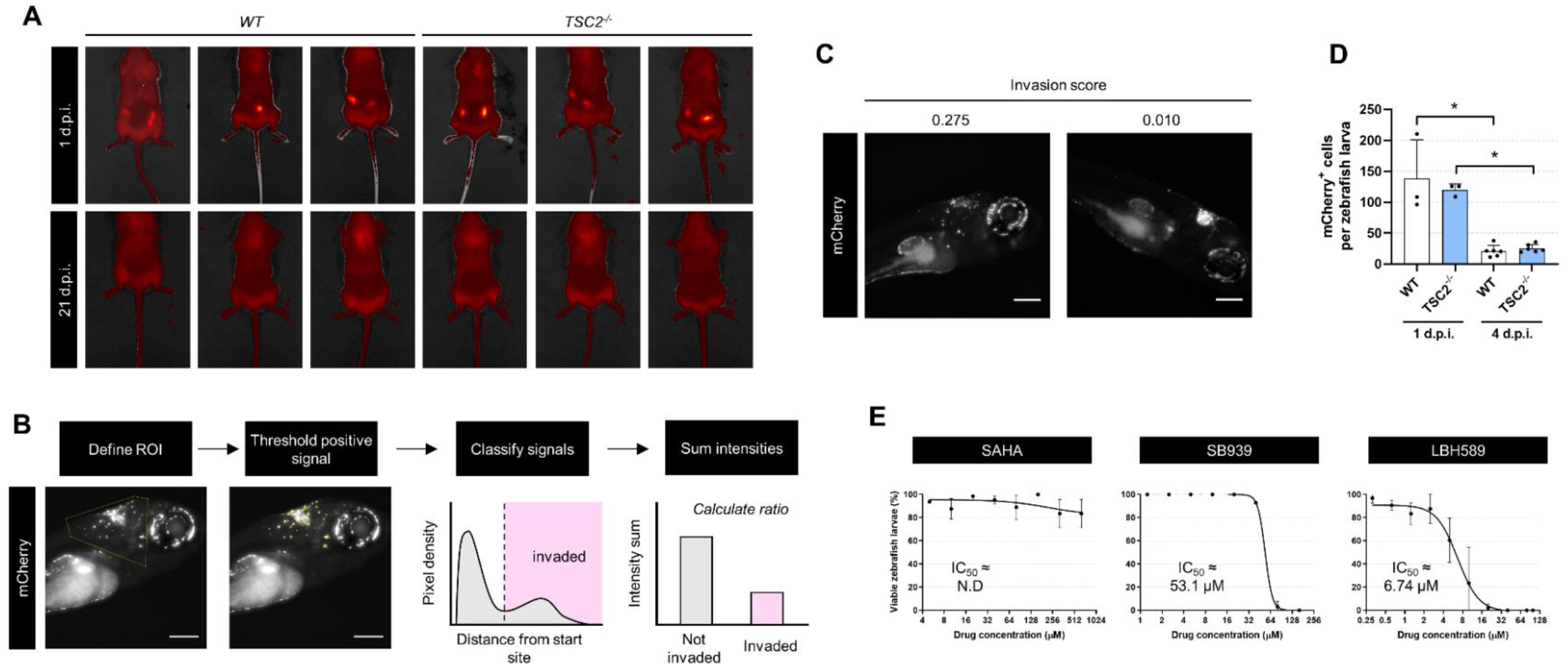
HDAC inhibitors are anti-invasive and selectively cytotoxic towards TSC2^-/-^ cells xenotransplanted into zebrafish. (A) Visualization of mCherry^+^ cells by IVIS following subcutaneous transplantation into rear flanks of immunodeficient NSG mice. (B) Schematic representation of invasion score calculation in zebrafish larvae. See Supplementary Materials and Methods for more details. Scale bars of 200µm. (C) Representative images of *TSC2^-/-^* mCherry^+^ cells disseminated 4 dpi with the associated invasion score. Scale bars of 200µm. (D) Number of mCherry^+^ cells per zebrafish detected by flow cytometry following whole larvae dissociation at 1 and 4 dpi. Each replicate is a pool of 15 – 20 zebrafish larvae (mean ± SD; * = p < 0.05 by Student t-test; n = 3 – 6). (E) Dose-toxicity curves of zebrafish larvae treated with HDAC inhibitors by immersion therapy.

### SUPPLEMENTARY TABLES

**Table S1.** Differential gene expression analysis of bulk RNA-seq data, untreated samples only. (A) DEG analysis of genotype (*TSC2^-/-^* vs. WT), controlling for substrate covariate. (B) DEG analysis of substrate (hydrogel vs. plastic), controlling for genotype covariate. (C) DEG analysis of interaction between genotype and ECM.

**Table S2.** GO term enrichment analysis of DEG lists. (A) GO term enrichment in *TSC2^-/-^* vs. WT DEG list (FDR < 0.05, |log_2_FC| > 1). (B) GO term enrichment in hydrogel vs. plastic DEG list (FDR < 0.05, |log_2_FC| > 1). (C) GO enrichment of genotype:substrate interaction DEG list (FDR < 0.05).

**Table S3.** Three-dimensional drug screen raw data. (A) Compound information from Ontario Institute of Cancer Research kinase inhibitor and tool compound libraries. (B-C) Cytotoxicity and invasion modulation effects of compounds, (B) statistic descriptions and (C) raw data.

**Table S4.** Enrichment results via adaptation of GSEA. Results for statistics of (A) selective cytotoxicity, positive enrichment (i.e., selectively cytotoxic towards *TSC2^-/-^*), (B) selective cytotoxicity, negative enrichment (i.e., selectively cytotoxic towards WT), (C) invasion modulation, positive enrichment (i.e., attenuate invasion), (D) invasion modulation, negative enrichment (i.e., potentiate invasion).

**Table S5.** Elion^TM^ structure-based compound analysis. (A-B) Significantly enriched targets and mechanisms of action by (A) selective cytotoxicity towards *TSC2^-/-^* cells and (B) invasion attenuation. (C-D) Significantly enriched GO and PFAM terms (based on significantly enriched targets) by (C) selective cytotoxicity towards *TSC2^-/-^* cells and (D) invasion attenuation.

### SUPPLEMENTARY MOVIES

**Movie S1.** Brightfield Z-stack of WT invading through the hydrogel, counterstained with Hoechst.

**Movie S2.** Brightfield Z-stack of *TSC2^-/-^* invading through the hydrogel, counterstained with Hoechst.

## Notes

### Competing Interest Statement

The authors have declared no competing interest.

## REFERENCES

1. A. M. Taveira-DaSilva, J. Moss, Clinical features, epidemiology, and therapy of lymphangioleiomyomatosis, Clin Epidemiol 7, 249–257 (2015).

2. J. Moss, N. A. Avila, P. M. Barnes, R. A. Litzenberger, J. Bechtle, P. G. Brooks, C. J. Hedin, S. Hunsberger, A. S. Kristof, Prevalence and Clinical Characteristics of Lymphangioleiomyomatosis (LAM) in Patients with Tuberous Sclerosis Complex, Am J Respir Crit Care Med 164, 669–671 (2001).

3. X. Zhe, L. Schuger, Combined Smooth Muscle and Melanocytic Differentiation in Lymphangioleiomyomatosis, J Histochem Cytochem. 52, 1537–1542 (2004).

4. G. F. Abbott, M. L. Rosado-de-Christenson, A. A. Frazier, T. J. Franks, R. D. Pugatch, J. R. Galvin, From the archives of the AFIP: lymphangioleiomyomatosis: radiologic-pathologic correlation, Radiographics 25, 803–828 (2005).

5. S. C. Chu, K. Horiba, J. Usuki, N. A. Avila, C. C. Chen, W. D. Travis, V. J. Ferrans, J. Moss, Comprehensive Evaluation of 35 Patients With Lymphangioleiomyomatosis, Chest 115, 1041–1052 (1999).

6. T. Urban, R. Lazor, J. Lacronique, M. Murris, S. Labrune, D. Valeyre, J. F. Cordier, Pulmonary lymphangioleiomyomatosis. A study of 69 patients. Groupe d’Etudes et de Recherche sur les Maladies “Orphelines” Pulmonaires (GERM“O”P)., Medicine (Baltimore) 78, 321–337 (1999).

7. E. P. Henske, F. X. McCormack, Lymphangioleiomyomatosis — a wolf in sheep’s clothing, J Clin Invest 122, 3807–3816 (2012).

8. A. M. Taveira-DaSilva, O. Hathaway, M. Stylianou, J. Moss, Changes in lung function and chylous effusions in patients with lymphangioleiomyomatosis treated with sirolimus, Ann. Intern. Med. 154, 797–805, W-292–293 (2011).

9. J. Bee, S. Fuller, S. Miller, S. R. Johnson, Lung function response and side effects to rapamycin for lymphangioleiomyomatosis: a prospective national cohort study, Thorax 73, 369–375 (2018).

10. J. Yao, A. M. Taveira-DaSilva, A. M. Jones, P. Julien-Williams, M. Stylianou, J. Moss, Sustained effects of sirolimus on lung function and cystic lung lesions in lymphangioleiomyomatosis, Am. J. Respir. Crit. Care Med. 190, 1273–1282 (2014).

11. F. X. McCormack, Y. Inoue, J. Moss, L. G. Singer, C. Strange, K. Nakata, A. F. Barker, J. T. Chapman, M. L. Brantly, J. M. Stocks, K. K. Brown, J. P. I. Lynch, H. J. Goldberg, L. R. Young, B. W. Kinder, G. P. Downey, E. J. Sullivan, T. V. Colby, R. T. McKay, M. M. Cohen, L. Korbee, A. M. Taveira-DaSilva, H.-S. Lee, J. P. Krischer, B. C. Trapnell, Efficacy and Safety of Sirolimus in Lymphangioleiomyomatosis. http://dx.doi.org/10.1056/NEJMoa1100391 (2011), doi:10.1056/NEJMoa1100391.

12. J. J. Bissler, F. X. McCormack, L. R. Young, J. M. Elwing, G. Chuck, J. M. Leonard, V. J. Schmithorst, T. Laor, A. S. Brody, J. Bean, S. Salisbury, D. N. Franz, Sirolimus for Angiomyolipoma in Tuberous Sclerosis Complex or Lymphangioleiomyomatosis, N Engl J Med 358, 140–151 (2008).

13. E. A. Goncharova, D. A. Goncharov, A. Eszterhas, D. S. Hunter, M. K. Glassberg, R. S. Yeung, C. L. Walker, D. Noonan, D. J. Kwiatkowski, M. M. Chou, R. A. Panettieri, V. P. Krymskaya, Tuberin regulates p70 S6 kinase activation and ribosomal protein S6 phosphorylation. A role for the TSC2 tumor suppressor gene in pulmonary lymphangioleiomyomatosis (LAM), J. Biol. Chem. 277, 30958–30967 (2002).

14. S. P. Delaney, L. M. Julian, A. Pietrobon, J. Yockell-Lelièvre, C. Doré, T. T. Wang, V. C. Doyon, A. Raymond, D. A. Patten, A. S. Kristof, M.-E. Harper, H. Sun, W. L. Stanford, Human pluripotent stem cell modeling of tuberous sclerosis complex reveals lineage-specific therapeutic vulnerabilities, bioRxiv, 683359 (2020).

15. D. J. Kwiatkowski, Animal Models of Lymphangioleiomyomatosis (LAM) and Tuberous Sclerosis Complex (TSC), Lymphatic Research and Biology 8, 51–57 (2010).

16. S. R. Caliari, J. A. Burdick, A practical guide to hydrogels for cell culture, Nature Methods 13, 405–414 (2016).

17. J. A. Burdick, G. D. Prestwich, Hyaluronic Acid Hydrogels for Biomedical Applications, Advanced Materials 23, H41–H56 (2011).

18. R. Y. Tam, J. Yockell-Lelièvre, L. J. Smith, L. M. Julian, A. E. G. Baker, C. Choey, M. S. Hasim, J. Dimitroulakos, W. L. Stanford, M. S. Shoichet, Rationally Designed 3D Hydrogels Model Invasive Lung Diseases Enabling High-Content Drug Screening, Advanced Materials 31, 1806214 (2019).

19. R. Edmondson, J. J. Broglie, A. F. Adcock, L. Yang, Three-dimensional cell culture systems and their applications in drug discovery and cell-based biosensors., Assay Drug Dev Technol 12, 207–218 (2014).

20. A. E. G. Baker, L. C. Bahlmann, R. Y. Tam, J. C. Liu, A. N. Ganesh, N. Mitrousis, R. Marcellus, M. Spears, J. M. S. Bartlett, D. W. Cescon, G. D. Bader, M. S. Shoichet, Benchmarking to the Gold Standard: Hyaluronan-Oxime Hydrogels Recapitulate Xenograft Models with In Vitro Breast Cancer Spheroid Culture, Advanced Materials 31, 1901166 (2019).

21. C. A. MacRae, R. T. Peterson, Zebrafish as tools for drug discovery, Nature Reviews Drug Discovery 14, 721–731 (2015).

22. D. C. Swinney, J. Anthony, How were new medicines discovered?, Nature Reviews Drug Discovery 10, 507–519 (2011).

23. L. M. Julian, S. P. Delaney, Y. Wang, A. A. Goldberg, C. Doré, J. Yockell-Lelièvre, R. Y. Tam, K. Giannikou, F. McMurray, M. S. Shoichet, M.-E. Harper, E. P. Henske, D. J. Kwiatkowski, T. N. Darling, J. Moss, A. S. Kristof, W. L. Stanford, Human Pluripotent Stem Cell–Derived TSC2-Haploinsufficient Smooth Muscle Cells Recapitulate Features of Lymphangioleiomyomatosis, Cancer Res (2017), doi:10.1158/0008-5472.CAN-17-0925.

24. M. Guo, J. J. Yu, A. K. Perl, K. A. Wikenheiser-Brokamp, M. Riccetti, E. Y. Zhang, P. Sudha, M. Adam, A. Potter, E. J. Kopras, K. Giannikou, S. S. Potter, S. Sherman, S. R. Hammes, D. J. Kwiatkowski, J. A. Whitsett, F. X. McCormack, Y. Xu, Single Cell Transcriptomic Analysis Identifies a Unique Pulmonary Lymphangioleiomyomatosis Cell, American Journal of Respiratory and Critical Care Medicine (2020), doi:10.1164/rccm.201912-2445OC.

25. H. Zhang, G. Cicchetti, H. Onda, H. B. Koon, K. Asrican, N. Bajraszewski, F. Vazquez, C. L. Carpenter, D. J. Kwiatkowski, Loss of Tsc1/Tsc2 activates mTOR and disrupts PI3K-Akt signaling through downregulation of PDGFR, J Clin Invest 112, 1223–1233 (2003).

26. A. Giese, M. A. Loo, N. Tran, D. Haskett, S. W. Coons, M. E. Berens, Dichotomy of astrocytoma migration and proliferation, Int. J. Cancer 67, 275–282 (1996).

27. A. J. Valvezan, B. D. Manning, Molecular logic of mTORC1 signalling as a metabolic rheostat, Nat Metab 1, 321–333 (2019).

28. D. S. Schrump, Cytotoxicity Mediated by Histone Deacetylase Inhibitors in Cancer Cells: Mechanisms and Potential Clinical Implications, Clin Cancer Res 15, 3947–3957 (2009).

29. A. S. Alzahrani, PI3K/Akt/mTOR inhibitors in cancer: At the bench and bedside, Seminars in Cancer Biology 59, 125–132 (2019).

30. A. L. Fridman, M. A. Tainsky, Critical pathways in cellular senescence and immortalization revealed by gene expression profiling, Oncogene 27, 5975–5987 (2008).

31. B. Adane, G. Alexe, B. K. A. Seong, D. Lu, E. E. Hwang, D. Hnisz, C. A. Lareau, L. Ross, S. Lin, F. S. Dela Cruz, M. Richardson, A. S. Weintraub, S. Wang, A. B. Iniguez, N. V. Dharia, A. S. Conway, A. L. Robichaud, B. Tanenbaum, J. M. Krill-Burger, F. Vazquez, M. Schenone, J. N. Berman, A. L. Kung, S. A. Carr, M. J. Aryee, R. A. Young, B. D. Crompton, K. Stegmaier, STAG2 loss rewires oncogenic and developmental programs to promote metastasis in Ewing sarcoma, Cancer Cell 39, 827–844.e10 (2021).

32. A. M. El-Naggar, C. J. Veinotte, H. Cheng, T. G. P. Grunewald, G. L. Negri, S. P. Somasekharan, D. P. Corkery, F. Tirode, J. Mathers, D. Khan, A. H. Kyle, J. H. Baker, N. E. LePard, S. McKinney, S. Hajee, M. Bosiljcic, G. Leprivier, C. E. Tognon, A. I. Minchinton, K. L. Bennewith, O. Delattre, Y. Wang, G. Dellaire, J. N. Berman, P. H. Sorensen, Translational Activation of HIF1α by YB-1 Promotes Sarcoma Metastasis, Cancer Cell 27, 682–697 (2015).

33. F. Yang, S. Sun, C. Wang, M. Haas, S. Yeo, J.-L. Guan, Targeted therapy for mTORC1-driven tumours through HDAC inhibition by exploiting innate vulnerability of mTORC1 hyper-activation, British Journal of Cancer, 1–12 (2020).

34. B. S. Mann, J. R. Johnson, M. H. Cohen, R. Justice, R. Pazdur, FDA approval summary: vorinostat for treatment of advanced primary cutaneous T-cell lymphoma, Oncologist 12, 1247–1252 (2007).

35. L. A. Raedler, Farydak (Panobinostat): First HDAC Inhibitor Approved for Patients with Relapsed Multiple Myeloma, Am Health Drug Benefits 9, 84–87 (2016).

36. T. E. Witzig, C. Reeder, J. J. Han, B. LaPlant, M. Stenson, H. W. Tun, W. Macon, S. M. Ansell, T. M. Habermann, D. J. Inwards, I. N. Micallef, P. B. Johnston, L. F. Porrata, J. P. Colgan, S. Markovic, G. S. Nowakowski, M. Gupta, The mTORC1 inhibitor everolimus has antitumor activity in vitro and produces tumor responses in patients with relapsed T-cell lymphoma, Blood 126, 328–335 (2015).

37. E. H. Rubin, N. G. B. Agrawal, E. J. Friedman, P. Scott, K. E. Mazina, L. Sun, L. Du, J. L. Ricker, S. R. Frankel, K. M. Gottesdiener, J. A. Wagner, M. Iwamoto, A Study to Determine the Effects of Food and Multiple Dosing on the Pharmacokinetics of Vorinostat Given Orally to Patients with Advanced Cancer, Clin Cancer Res 12, 7039–7045 (2006).

38. F. Giles, T. Fischer, J. Cortes, G. Garcia-Manero, J. Beck, F. Ravandi, E. Masson, P. Rae, G. Laird, S. Sharma, H. Kantarjian, M. Dugan, M. Albitar, K. Bhalla, A Phase I Study of Intravenous LBH589, a Novel Cinnamic Hydroxamic Acid Analogue Histone Deacetylase Inhibitor, in Patients with Refractory Hematologic Malignancies, Clin Cancer Res 12, 4628– 4635 (2006).

39. W. P. Yong, B. C. Goh, R. A. Soo, H. C. Toh, K. Ethirajulu, J. Wood, V. Novotny-Diermayr, S. C. Lee, W. L. Yeo, D. Chan, D. Lim, E. Seah, R. Lim, J. Zhu, Phase I and pharmacodynamic study of an orally administered novel inhibitor of histone deacetylases, SB939, in patients with refractory solid malignancies, Ann Oncol 22, 2516–2522 (2011).

40. C. Zhang, V. Richon, X. Ni, R. Talpur, M. Duvic, Selective Induction of Apoptosis by Histone Deacetylase Inhibitor SAHA in Cutaneous T-Cell Lymphoma Cells: Relevance to Mechanism of Therapeutic Action, Journal of Investigative Dermatology 125, 1045–1052 (2005).

41. R. M. White, A. Sessa, C. Burke, T. Bowman, J. LeBlanc, C. Ceol, C. Bourque, M. Dovey, W. Goessling, C. E. Burns, L. I. Zon, Transparent Adult Zebrafish as a Tool for In Vivo Transplantation Analysis, Cell Stem Cell 2, 183–189 (2008).

42. M. Westerfield, THE Zebrafish Book a Guide for the Laboratory Use of Zebrafish Danio*.

43. A. Subramanian, P. Tamayo, V. K. Mootha, S. Mukherjee, B. L. Ebert, M. A. Gillette, A. Paulovich, S. L. Pomeroy, T. R. Golub, E. S. Lander, J. P. Mesirov, Gene set enrichment analysis: A knowledge-based approach for interpreting genome-wide expression profiles, Proc. Natl. Acad. Sci. 102, 15545–15550 (2005).

44. D. P. Corkery, G. Dellaire, J. N. Berman, Leukaemia xenotransplantation in zebrafish – chemotherapy response assay in vivo, Br. J. Haematol. 153, 786–789 (2011).

45. H. M, T. C, S. Wl, M. P, Human melanoma cells transplanted into zebrafish proliferate, migrate, produce melanin, form masses and stimulate angiogenesis in zebrafish., Angiogenesis 9, 139–151 (2006).

